# Glycoproteomics-compatible MS/MS-based quantification of glycopeptide isomers

**DOI:** 10.1101/2023.01.31.526390

**Authors:** Joshua C.L. Maliepaard, J. Mirjam A. Damen, Geert-Jan P.H. Boons, Karli R. Reiding

**Affiliations:** Biomolecular Mass Spectrometry and Proteomics, Utrecht Institute for Pharmaceutical Sciences and Bijvoet Center for Biomolecular Research, University of Utrecht, Utrecht, the Netherlands; Netherlands Proteomics Center, Utrecht, the Netherlands; Department of Chemical Biology and Drug Discovery, Utrecht Institute for Pharmaceutical Sciences and Bijvoet Center for Biomolecular Research, University of Utrecht, Utrecht, the Netherlands; Complex Carbohydrate Research Center, University of Georgia, Athens, GA, USA; Department of Biochemistry and Molecular Biology, University of Georgia, Athens, Ga, USA

**Keywords:** glycosylation, glycopeptide, isomers, isomerism, linkage, sialylation, galactosylation, stepped HCD

## Abstract

Glycosylation is an essential protein modification occurring on the majority of extracellular human proteins, mass spectrometry (MS) being an indispensable tool for its analysis. Not only can MS determine glycan compositions, but also position the glycan at specific sites via glycoproteomics. However, glycans are complex branching structures with monosaccharides interconnected in a variety of biologically relevant linkages - isomeric properties which are invisible when the readout is mass alone.

Here, we developed an LC-MS/MS-based workflow for determining glycopeptide isomer ratios. Making use of isomerically-defined glyco(peptide) standards, we observed marked differences in fragmentation behavior between isomer pairs when subjected to collision energy gradients, specifically in terms of galactosylation/sialylation branching and linkage. These behaviors were developed into component variables that allowed relative quantification of isomerism within mixtures. Importantly, at least for small peptides, the isomer quantification appeared largely independent from the peptide portion of the conjugate, allowing broad application of the method.

## INTRODUCTION

Glycosylation is a ubiquitous form of post-translational modification (PTM) that occurs on the majority of extracellular proteins(1,2). Protein *N*-glycosylation, glycosylation of the side-chain of an Asn residue, primarily takes place within an Asn-Xxx-Ser/Thr motif (Xxx ≠ Pro) and often on multiple sites within the same protein(2). Glycans themselves consist of monosaccharides interconnected via glycosidic bonds; typical monosaccharides include galactose (Gal), mannose (Man) and glucose (Glc), collectively referred to as hexoses (Hex) due to their shared six-ring and mass, *N*-acetylglucosamine (GlcNAc) and *N*-acetylgalactosamine (GalNAc), referred to as *N*-acetylhexosamines (HexNAc), all in addition to fucoses (Fuc) and sialic acids of which *N*-acetylneuraminic acid (NeuAc) is the primary human variant(3). The carbon with which one monosaccharide connects to another determines the terminology of the glycan linkage; for each pair of monosaccharides, a set of numbers describes which carbons are joined via *O*-glycosidic linkage, *e*.*g*., 1-4, whereas the addition of the prefix α or β informs on the anomerism of the bond(1). A typical example of this is lactose, in which a Gal is β1-4-linked to a Glc, *i*.*e*., Galβ1-4Glc(1). The wealth of possible linkages between monosaccharides, and the resulting branching glycan structures, represents a large reservoir of biological variation between one glycoproteoform and the next.

Glycans are involved in a variety of cellular processes and even a slight change in the structure of a glycan can have a major impact on the function of a glycoprotein or the way it interacts with its environment. One example is the way in which sialic acid linkage effects the ability of influenza viruses like the H5N1 avian influenza to cross the species barrier(4). Avian influenza viruses mostly infect via binding α2,3-linked sialylation, which lines the avian respiratory track, while the human tract is mostly lined with α2,6-linked sialylation(5). However, swine carry both α2,3- and α2,6-linked sialic acids, making them ideal candidates for cross-species contamination and subsequent mutation(5). When treating a mouse arthritis model with intravenous immunoglobulin, α2,6-immunoglobulin G (IgG) was reported to improve the anti-inflammatory effect over α2,3-linked sialylation(4). In terms of glycan branching, monogalactosylation of the fragment crystallizable (Fc) domain of IgG glycan was linked to higher complement-mediated toxicity (CDC) activity(6). This importance of glycan structure for antibody activity combined with the rise use of antibodies in therapeutics – 9 out of the 20 top selling therapeutics in the USA in 2019 were antibodies(7) – deciphering glycopeptide structure is more important than ever.

Mass spectrometry (MS) is a powerful tool to perform glycosylation analysis – the mass of a glycan, glycopeptide or glycoprotein is highly informative for acquiring the composition of glycosylation(8). However, glycan isomerism is extremely challenging to study from mass alone. For released glycan analysis, certain isomeric aspects of glycosylation can already be determined by the separation method prior to the MS analysis. For instance, porous graphitized carbon liquid chromatography (PGC-LC) can separate α2,3-/α2,6-sialylation and β1,3-/β1,4-galactosylation(9,10), hydrophilic interaction liquid chromatography (HILIC) allows distinction between sialic acid linkages and galactose branching(11,12), while capillary electrophoresis has been instrumental in separating the sialic acid linkages as well(13). Next to this, chemical derivatization of sialic acids is an attractive strategy to add a mass tag to different linkages(14), collision-induced dissociation can provide isomerism-dependent fragments and fragment ratios(15,16), and ion mobility (IM)-MS is rapidly maturing as a gas-phase separation method for analyzing isomerism as well(17–19). However, despite the extensive development in determining the isomerism of released glycans, work on glycopeptides remains very sparse. This class of analytes can provide site-specific information - essential for multiply-glycosylated proteins or protein mixtures - but the peptide portion presents a major analytical challenge. Sialic acid acid linkages appear accessible from an MS^2^ or MS^3^ level, although its assessment was primarily performed for *O*-glycopeptides(16). As it is, there currently appear to be no proteomics-compatible methods to determine glycan structural isomerism, let alone the ratios thereof for a given glycosylation site.

As such, we here present a proteomics-compatible method for the relative quantification of isomer ratios at the glycopeptide level, thus far suitable for the determination of sialic acid linkages and sialic acid/galactose branching positions. We have developed this method by using isomerically defined glycosylated asparagine standards, generated by chemoenzymatic synthesis(20), and show broad applicability by analyzing isomer mixtures and differing peptide lengths. Importantly, the determination of glycopeptide isomerism only requires MS/MS, namely a cycle through collision energies, and can therefore easily be applied without using special instrumentation or modes of fragmentation.

## RESULTS

### Standard identities

For MS determination of isomerism, we made use of eight isomerically-defined glycosylated asparagine standards, as synthesized and characterized previously(21,22). Combined, these standards covered four distinct isomeric properties (**Figure 1**); three out of the four pairs covered branching, *i*.*e*., having antennary extension on either the α1,3- or α1,6-linked Man allowing the investigation of Gal-, NeuAcGal and NeuAc-branching, whereas the fourth isomer pair differed in linkage of the NeuAc, *i*.*e*., full occupation of either α2,3- or α2,6-linked NeuAc. Accordingly, the respective compositions of the standards were N4H4, N4H4S1, N4H5S1 and N4H5S2 (N = *N*-acetylhexosamine, H = hexose, S = *N*-acetylneuraminic acid).

**Figure 1.**
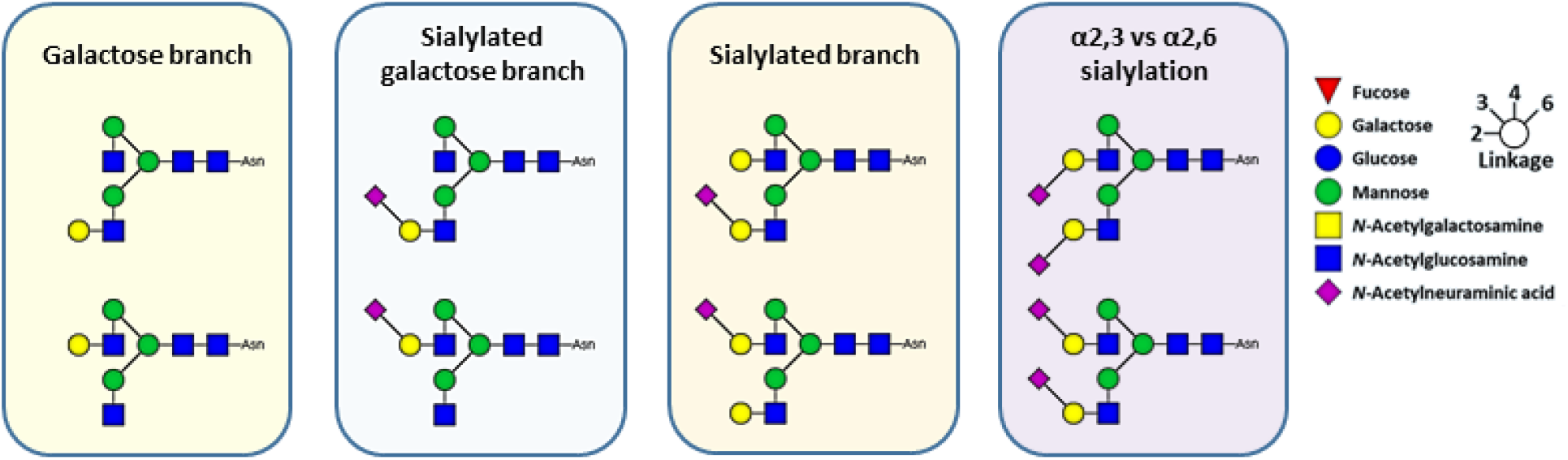
Overview of isomerically-defined glycosylated asparagine standards. A total of eight isomerically defined glycopeptides were combined into four pairs of linkage characteristics. Three pairs covered branching position, having antennary extension on either the α1,3- or α1,6-linked Man, allowing the investigation of Gal-, NeuAcGal and NeuAc-branching. The other glycopeptide pair differed in sialic acid linkage, allowing the investigation of α2,3- and α2,6-sialylation.

### Fragmentation behavior across NCE range

The isomerically defined glycosylated asparagine standards were analyzed by LC-MS/MS-based glycoproteomics with varying normalized collision energies (NCEs). Expectedly, the chromatography of the isomer pairs exhibited very similar peak shapes and retention times, alongside an identical number of isotopes and distribution thereof in MS (**Figure 2A,B**). At the same time, clear differences in oxonium ion intensities could be observed between the isomer pairs when subjected to MS/MS at given NCEs, for example between α2,3- and α2,6-sialylated glycosylated asparagine standards at 20% NCE (**Figure 2C**). Here, the relative abundances for *N*-acetylneuraminic acid (NeuAc) oxonium ions with and without water loss (*m/z* 274.0921 and 292.1027) both proved approximately 40% of the base peak intensity (BPI) for the α2,3-linked variant, while these were only 15 and 20% for the α2,6-linked variant. Similarly, the oxonium ions for HexNAc (*m/z* 204.0866) and HexHexNAc (*m/z* 366.1395) showed BPIs of 58 and 68% for the α2,3-sialylated glycopeptide standard and 54 and 100% for its α2,6-sialylated equivalent. The differences in MS/MS were not constrained to the sialic acid linkages, but could be observed for branching positions as well. For example, at 10% NCE, 3-branch monogalactosylated glycopeptides showed for HexNAc and HexHexNAc ions abundances of 44 and 100% BPI, respectively, whereas the variant with the galactose on the 6-branch showed abundances of 100 and 28% instead.

**Figure 2:**
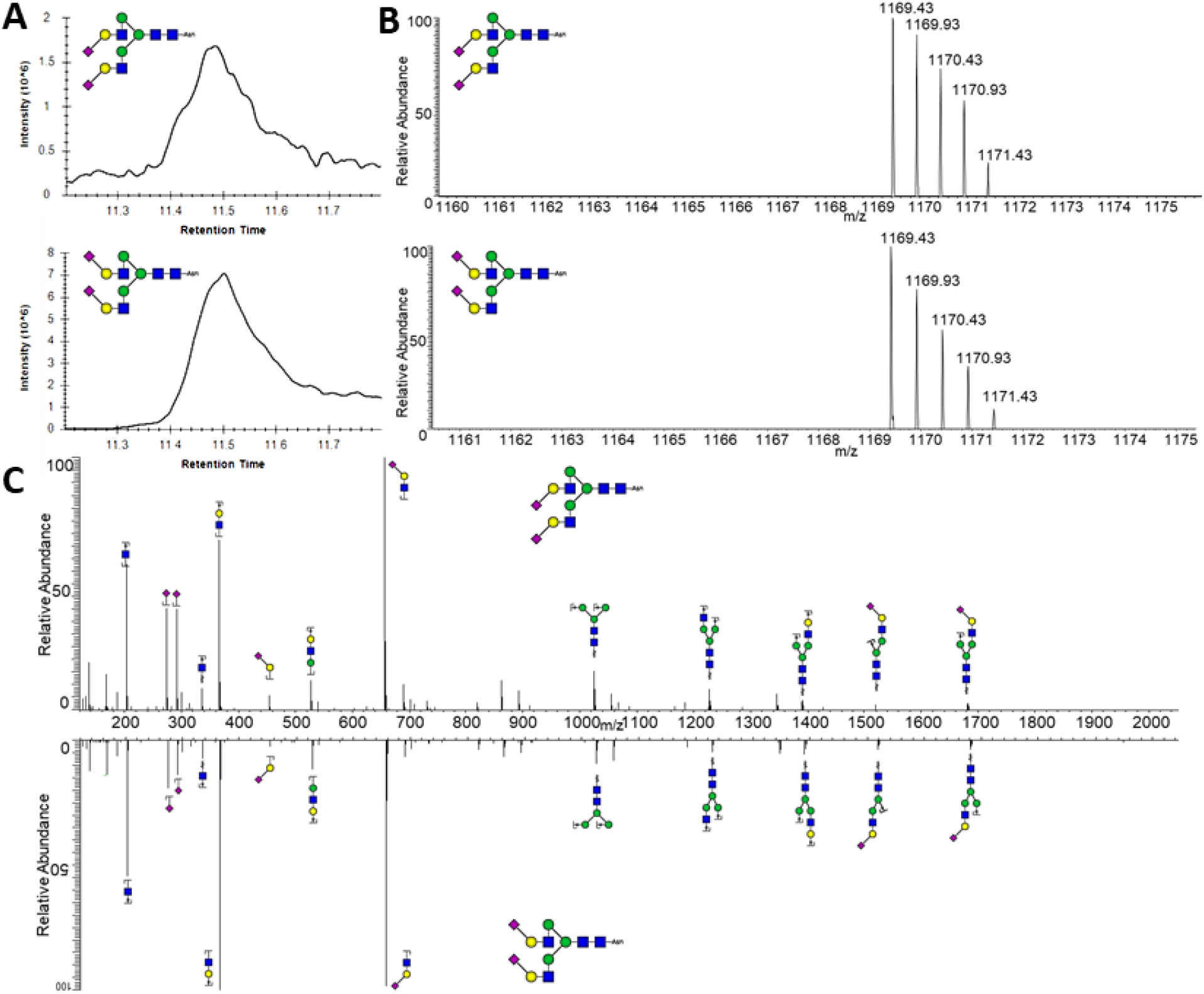
Comparison of N4H5S2 isomers in LC, MS and MS/MS. **A)** Separation of sialic acid linkage isomers by C18 chromatography. **B**) 2+ signals of the sialic acid linkage isomers in MS. C) MS/MS spectra generated by the sialic acid linkage isomers when subjected to HCD at NCE 20%. As can be seen, while LC and MS do not show differences between the isomers, MS/MS shows discernible peak ratio differences.

These profound differences led us to investigate the behavior of a wider group of ions across the collision energy space (**Figure 3**). In order to maximize the amount of MS/MS scans of glycopeptide, a mass trigger was introduced: when at least 3 of oxonium ions from a list of 15 were detected in a preliminary MS/MS scan at 29% NCE, 10 subsequent MS/MS scans of the same precursor were performed at NCEs ranging from 5-50%. The NCE range was set at 5-50% NCE since NCE below 5% showed almost no fragmentation of the parent ion, while at NCE 50% the spectrum was dominated by *m/z* 138.0550 with very few other oxonium ions present. A step size of 5% NCE was chosen in order to keep the total scan cycle time in line with other proteomics methods (∼250 ms) while still giving a good coverage of the entire NCE range. Obtained intensities of the oxonium ions were normalized to the total intensity, and compared between isomer pairs (**Figure 4**). Only oxonium ions (B-ions) were taken into account, as opposed to Y-ions, since these can principally be formed irrespective of the connecting peptide thereby making translation to alternative peptide backbones more likely.

**Figure 3.**
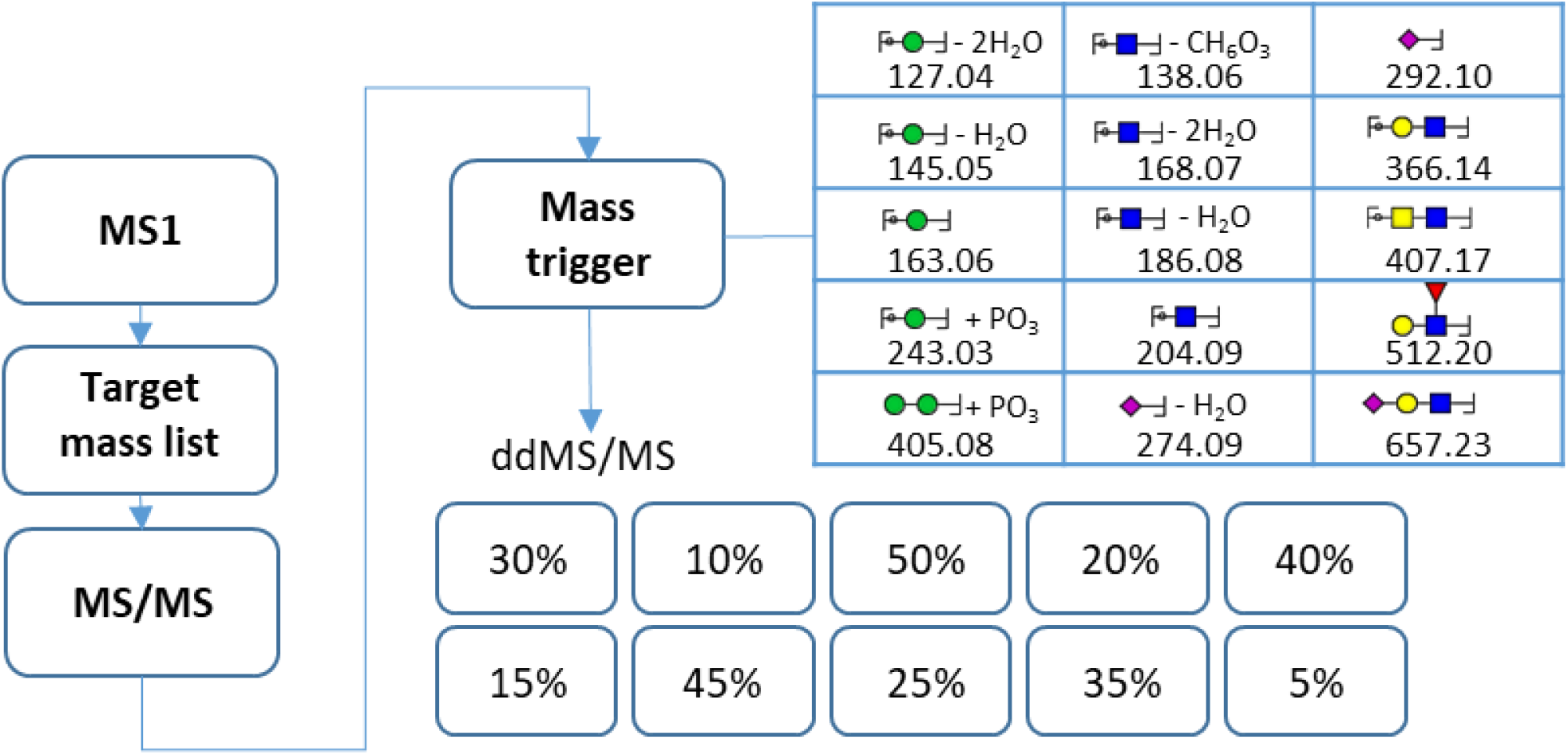
Stepped HCD LC-MS/MS method employed throughout the experiments. Following MS1, the LC-MS/MS method selects precursors from a target mass list, covering, depending on the application, isomeric standards, sialylglycopeptides and trastuzumab glycopeptides. Hereafter an MS/MS prescan is performed at HCD NCE 29%. If this prescan contains at least 3 detections matching a mass trigger list with glycan-specific oxonium ions, a series of stepped HCD MS/MS scans is triggered covering NCEs 5-50% in steps of 5%, following a semi-randomized fashion.

**Figure 4:**
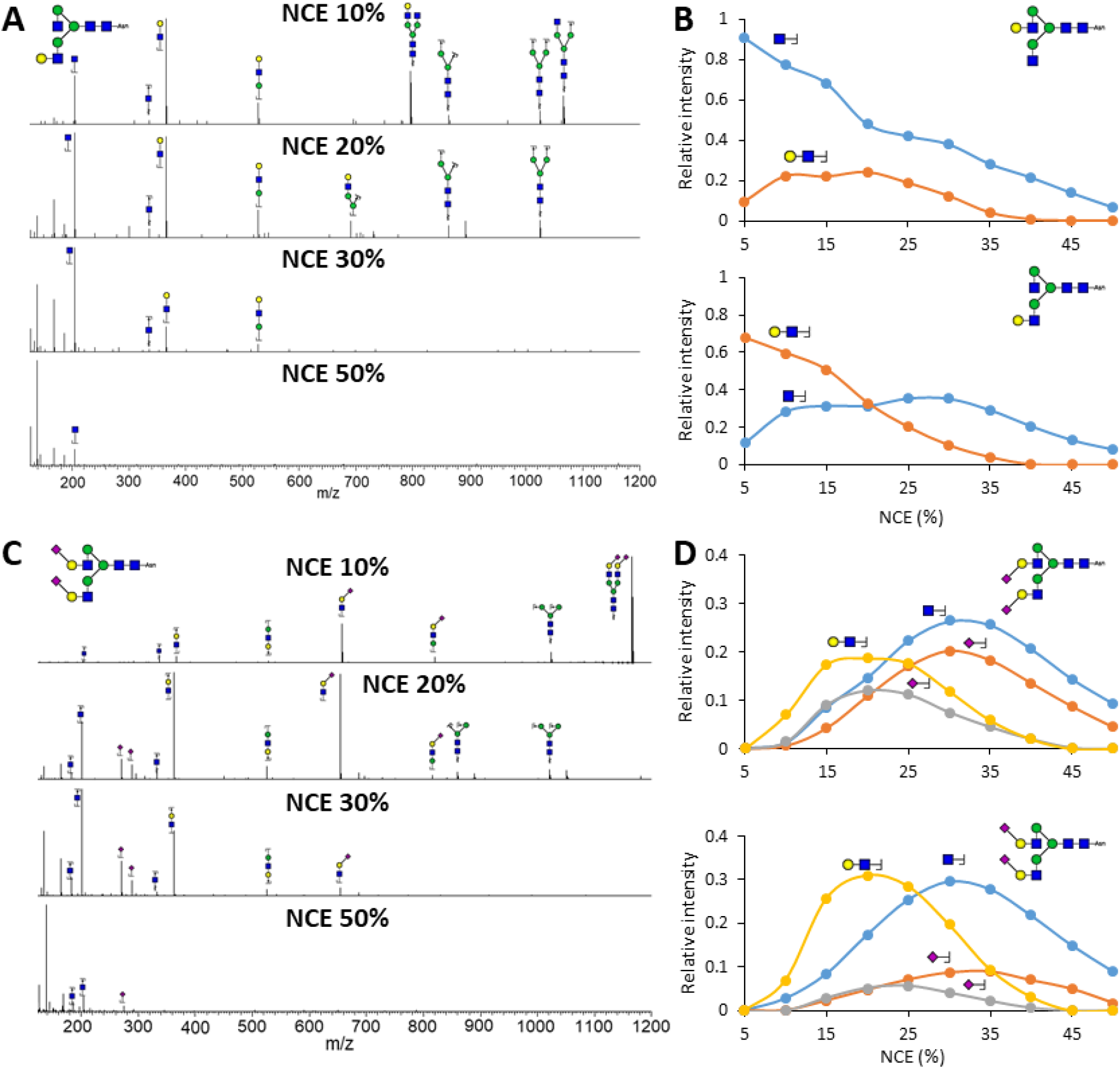
MS/MS differences between glycopeptide isomers. **A**) MS/MS at NCEs of 10, 20, 30 and 50% of the N4H4 glycan with galactosylation positioned on the 6-branch. **B**) Relative intensity differences between the oxonium ions HexNAc and HexHexNAc for 5-50% NCE, compared between isomers with the galactose on the 6-branch (**top**) and 3-branch (**bottom**). **C**) MS/MS spectra at NCEs of 10, 20, 30 and 50% of a fully α2,6-sialylated N4H5S2 glycan. **D**) Relative intensity differences of the oxonium ions HexNAc, HexHexNAc, NeuAc-H_2_O and NeuAc for 5-50% NCE, compared between sialylation isomers with full α2,3-linkage (**top**) and α2,6-linkage (**bottom**). For both the galactosylation and sialylation isomers, NCE regions could be determined in which the isomers were distinguishable.

To determine which charge states would be most beneficial for distinguishing the glycopeptide isomers, the standard panel was primarily measured with a stepped-HCD method that allowed for fragmentation of analyte charge states ranging from 2+ to 8+. Interestingly, the main charge states observed for the standards, namely 2+ and 3+, showed noticeable fragmentation differences, with 2+ having much more pronounced relative intensity differences between isomer pairs than 3+ (**Figure 5**). One example of this was the relative intensity of HexNAc (*m/z* 204.0867), which 10% NCE in 2+ was 77% BPI for 6-branched galactosylation and 28% BPI for its isomeric counterpart (a 49% difference), whereas in 3+ these values were 48% BPI and 43% BPI (a 5% difference). For NeuAc-H_2_O (*m/z* 274.0921) at 30% NCE in 2+, the relative intensity was 20% BPI for α2,3-linked sialylation and 9% BPI for its isomer (a 11% difference), whereas in 3+ these numbers were 22% BPI and 15% BPI (a 7% difference) (**Figure 5**). Because of this, the decision was made to proceed with measurement and data analysis of analytes with charge state 2+ specifically.

**Figure 5:**
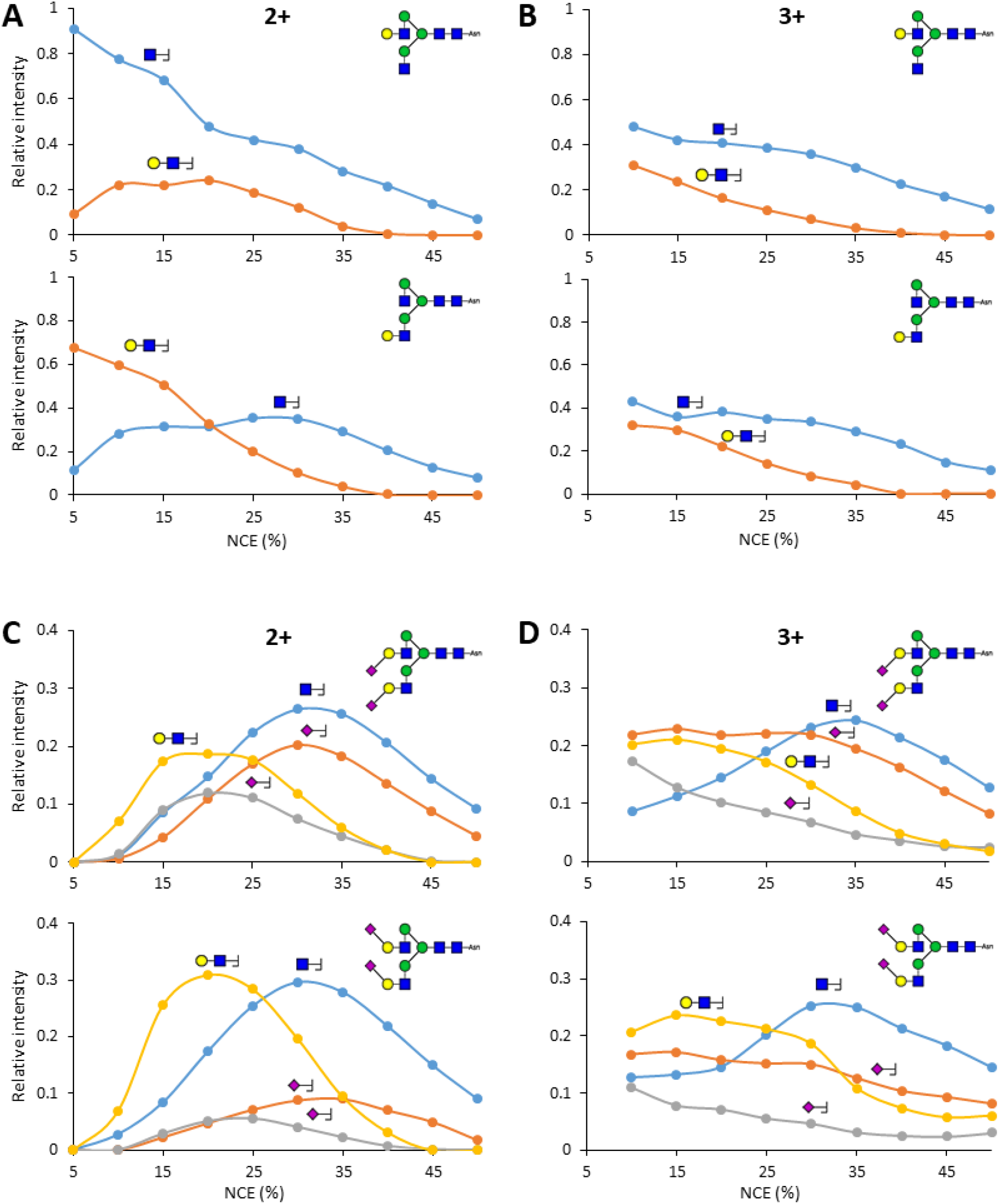
MS/MS fragmentation changes for charge states 2+ and 3+. **A**) Relative intensity differences between the oxonium ions for HexNAc and HexHexNAc, compared between galactose isomers, for precursor charge states 2+ (**left**) and 3+ (**right**). **B**) Relative intensity differences between the oxonium ions for HexNAc, HexHexNAc, NeuAc-H_2_O and NeuAc, compared between sialic acid linkage isomers, for precursor charge states 2+ (left) and 3+ (right). As can be seen, the ion ratio differences between isomers are more pronounced in 2+ when compared to 3+.

### Construction of variable for isomer determination

Since clear differences were observed in the fragmentation behavior of the isomeric standards across collision energy, the next step was to attempt relative quantification within mixtures thereof. For this purpose, isomerically defined glycosylated asparagine standards were mixed ranging from 100% of one isomer (A) to 100% of the other (B) in steps of 10%. After LC-MS measurement, for the relative intensity for 1) each oxonium ion at 2) each measured collision energy across 3) each glycopeptide mixture, was interpreted for the purpose of isomer discrimination (**Figure 6, Supplementary Figures 1-4**).

**Figure 6:**
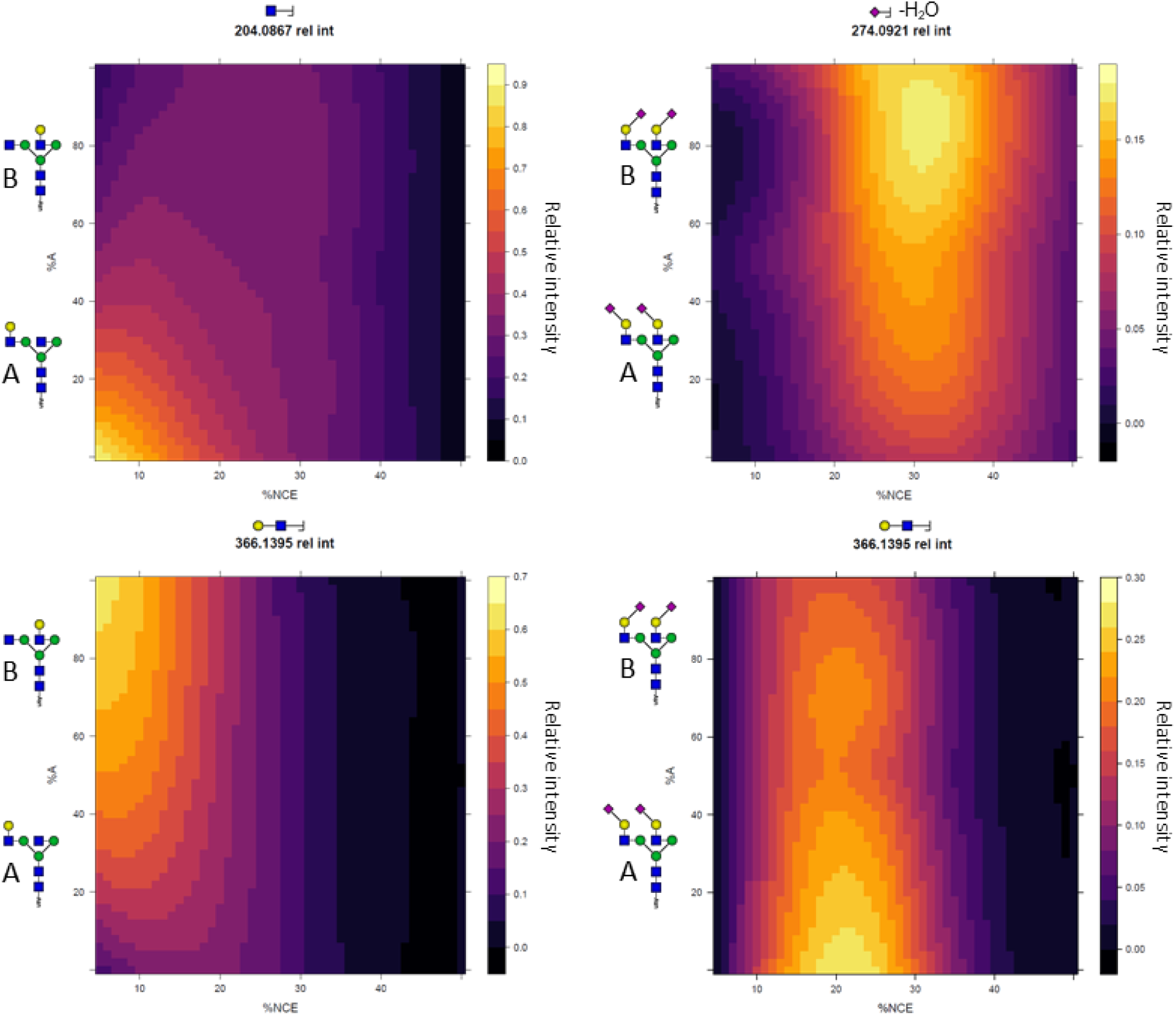
Oxonium ion intensity differences across isomer mixtures. Each panel represents a single oxonium ion. Displayed on the x-axis is the collision energy gradient in % NCE, on the y-axis the ratio of the isomer pair in %A (left isomer), and on the z-axis (the color) the intensity of the oxonium ion, normalized against all others within the same MS-run. As can be seen, not only do the presented oxonium ions showed differences between isomers across the energy gradient, but they gradually changed across the isomer mixture as well.

Taking the monogalactosylated glycopeptide mixtures as an example, at an NCE of 10%, the relative abundance of the HexNAc oxonium ion (*m/z* 204.0867) increased from 20% base peak intensity (BPI) in the sample with the galactose on the 3-branch, to 80% BPI in the samples with the galactose on the 6-branch. At the same time, the relative abundance of the HexHexNAc ion (*m/z* 366.1395) increased from 25% to 60% BPI. In general, the differences in relative oxonium ion abundance were clearest at collision energies ranging from 10-25% NCE. For the α2,3- and α2,6-linked sialylation, the main MS/MS intensity differences proved to be NeuAc (*m/z* 292.1027) and NeuAc-H_2_O (*m/z* 274.0921), most distinguishable between NCEs of 20% to 45%. For example, at NCE 30% the relative intensity of NeuAc-H_2_O moved from 9% to 18% BPI when changing linkage from α2,6 to α2,3, while at the same time NeuAc increased from 5% to 10% (NCE 20%). Other oxonium ions most likely derived from the glycan branches, such as HexHexNAc (*m/z* 366.1395) and HexNAc (*m/z* 204.0867), showed instead the highest relative abundance with full α2,6-sialylation, gradually decreasing with increasing α2,3-linkage.

Since the differences in HCD fragmentation between isomers occurred over a range of collision energies and for multiple oxonium ions, a variable was created to condense these aspects into a single number. For the monogalactosylated glycopeptides, the oxonium ions that showed the greatest difference were HexNAc (*m/*z 204.0867) and HexHexNAc (*m/*z 366.1395), most appreciably so between NCE 10 and 25%. Below NCE 10% the ion intensity appeared to be too variable, whereas above 25% the isomers no longer showed a difference. For variable construction, then, the relative abundance of HexNAc (*m/z* 204.0867) was divided by the sum of the relative abundance of the HexHexNAc and HexNAc (*m/z* 366.1395 and 204.0867, respectively), averaged across NCE 10-25% (*i*.*e*., divided by the number of included collision energies to yield a number between zero and one). The same process was followed for the sialylated and sialylated galactose glycopeptide isomers: for N4H4S1 both HexHexNAc and HexHexNAcNeuAc (respectively *m/z* 366.1395 and 657.2349) showed the largest difference across NCEs 10-15%, making the formula for linkage determination 366.1395 / (366.1395 + 657.2349) / 2, whereas for N4H5S1 the ions HexNAc (*m/z* 204.0867), HexHexNAc (*m/z* 366.1395) and HexHexNAcNeuAc (*m/z* 657.2349) led up to a linkage variable formula of 204.0867 / (204.0867 + 366.1395 + 657.2349) / 3 between NCEs 10-20%. Finally, the difference in α2,3 and α2,6-sialylation for N4H5S2 proved most determinable by (274.0921 + 292.1027) / (274.0921 + 292.1027 + 204.0921 + 366.1395) averaged across NCE 20-45%.

Across the isomer mixtures, the linkage variables showed clear statistically significant differences between mixture percentage steps (**Figure 7**). For example, a two-tailed students t-test showed a *p*-value < 0.05 for 0-10%, 10-20%, 20-30%, 40-50% and 90-100% α2,3-linkage with 1.64·10^−7^ when comparing 0 and 10% α2,3-linkage, and 0.048 when comparing 20% and 30% α2,3-linkage as examples. Also comparisons for galactose branching, with the exception of 60-70% and 90-100%, consistently showed p-values below 0.01, examples being *p* = 1.99·10^−6^ for 30-40% and *p* = 4.64·10^−4^ for 70-80%. This allowed us to quantify the glycan structures of most of the tested glycopeptide mixtures with an approximate accuracy of 10%

**Figure 7:**
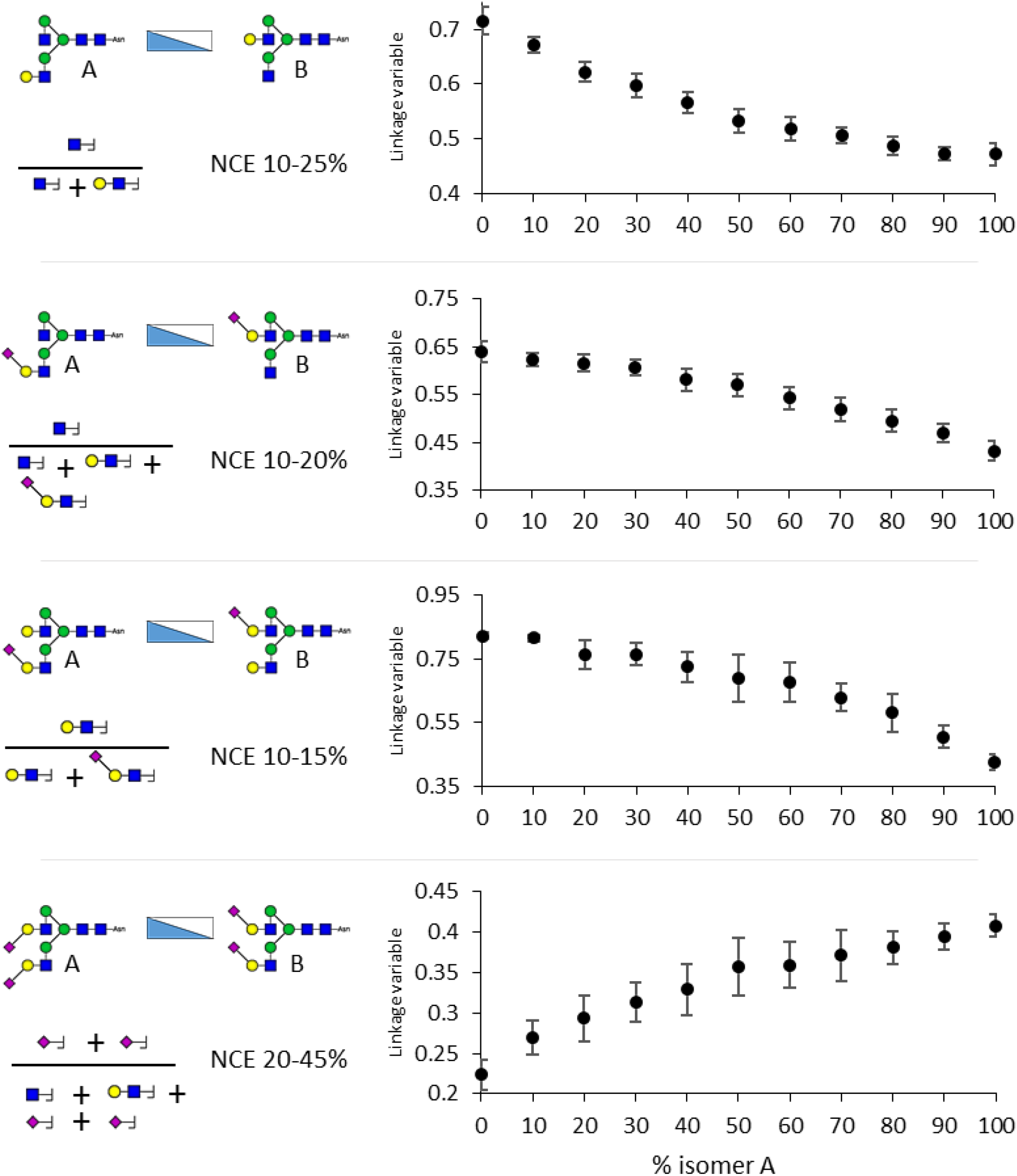
Average linkage variables for glycopeptide isomer mixes with standard deviations. The linkage variables for the glycopeptide isomers were based on the relative intensity of glycan oxonium ions across a range of collision energies, as indicated on the left. Already at differences of 10%, most mixtures differed with statistical significance.

To further ensure data quality and remove obvious outliers, the following curation criteria were used: A 10-20 min retention time window for samples run with a 60 min gradient, a 5-20 min retention time window for samples run with a 30 min gradient, a 5·10^3^ intensity threshold for NeuAc-H_2_O (*m/z* 274.0921) and a 1·10^4^ intensity threshold for HexNAc, HexHexNAc, and NeuAcHexHexNAc (*m/z* 204.0867, 366.1395, 657.2349 respectively) in the samples in which they occurred. This data curation led to the elimination of MS/MS spectra that would otherwise be wrongfully included in the variable, such as apparent alternative glycopeptide compositions (**Supplementary Figure 5**).

### Glycopeptide analysis

Subsequently, the effect of peptide backbone on the linkage variable was tested to determine the applicability of the linkage variables for glycopeptides with different peptide backbones but the same glycan.

Sialylglycopeptide (SGP), a well-studied glycopeptide with a known glycan structure(23), was used to assess whether the length of the connected peptide impacted fragmentation characteristics, and whether this could be overcome by the linkage variable. To obtain a range of different peptides with the same glycan, SGP was partially digested with Pronase. This yielded a range of glycopeptides of different sizes, *i*.*e*., N (identical to the standard panel), AN, NKT, KVAN/VANK (same mass), VANKT and KVANKT (the full peptide) (**Supplementary Figure 6**). Using the same LC-MS/MS method as for the isomerically defined glycosylated asparagine standards, the linkage variables were determined for the different glycopeptides carrying a N4H5S2 glycan. The linkage variable ranged from 0.20 to 0.25 across the glycopeptides, with the linkage variable of NKT being highest at 0.25 on average, and lowest being NK at 17%. All other peptides showed linkage variables between 0.21 and 0.24 (**Figure 8A**). These linkage variables corresponded with almost exclusively (90-100%) α2,6-linked sialic acids, which was in accordance with the literature(23). The similarity of the linkage variables indicated that the presence of the amino acids A, K, V and T did not directly affect the linkage variable to a large degree.

**Figure 8:**
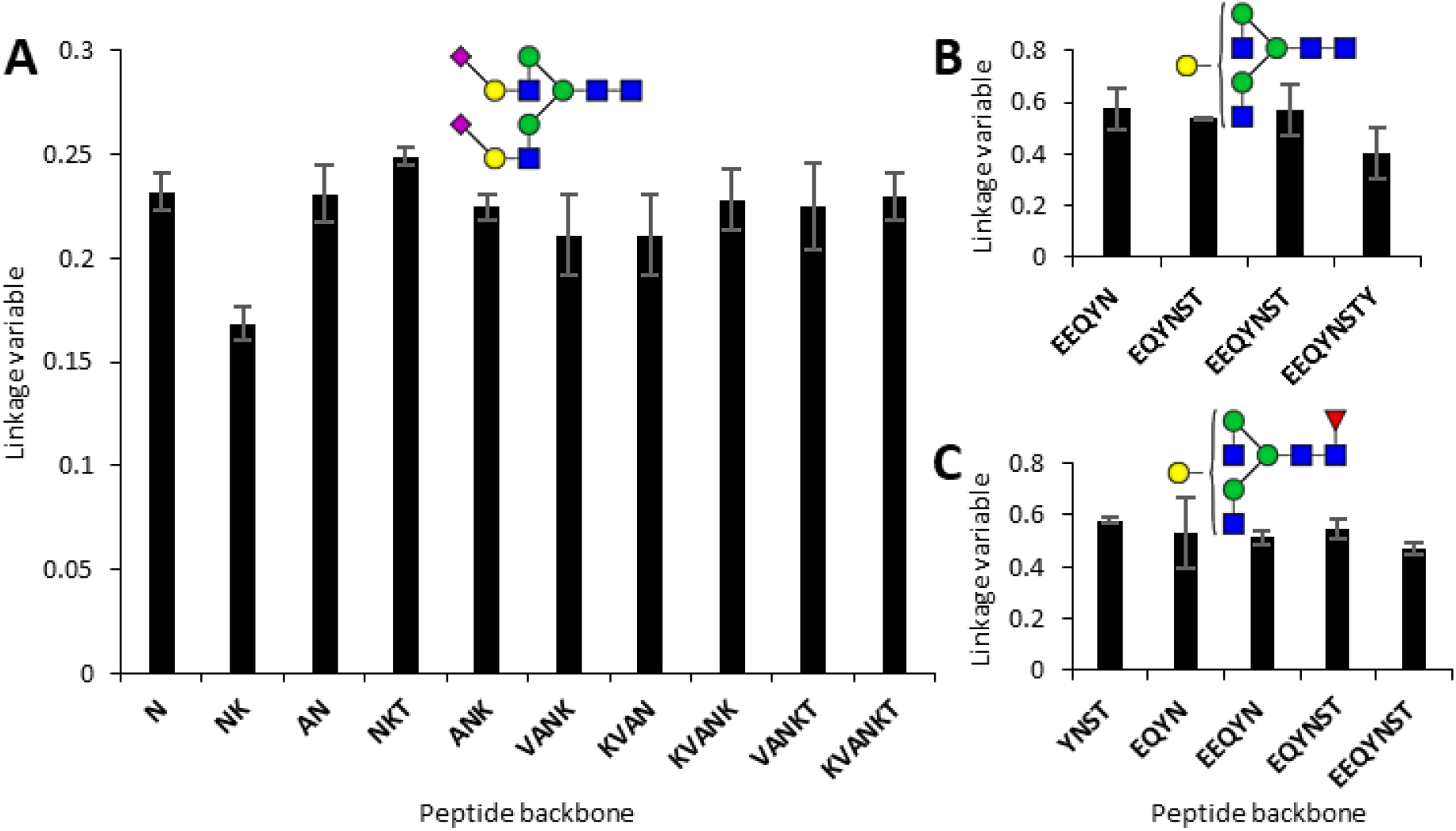
Effect of peptide backbone on sialic acid linkage and galactose branching assignment. **A)** Reported here are average linkage variables ± standard deviation for partially digested SGP with N4H5S2. As can be seen, the linkage variable ranged from 0.2 to 0.25 across the glycopeptides, with the average value of NKT being highest at 0.25 and lowest being NK at 0.17. These linkage variables corresponded with almost exclusively (90-100%) α2,6-linked sialic acids, which was in accordance with the literature for SGP(23). **B)** Reported here are average linkage variables ± standard deviation for partially digested trastuzumab with N4H4. As can be seen, the linkage variable ranged from 0.4 to 0.57 across the glycopeptides, with an average value of 0.54. This linkage variable corresponds to an approximate of 40-50% galactosylation of the 3-branch **C)** Reported here are average linkage variables ± standard deviation for partially digested trastuzumab with N4H4F1. As can be seen, the linkage variable ranged from 0.47 to 0.58 across the glycopeptides, with an average value of 0.51. This linkage variable corresponds to an approximate of 60-70% galactosylation of the 3-branch.

Next to SGP, trastuzumab was chosen as a glycoprotein to assess the effect of the peptide backbone on the linkage variable. Trastuzumab, also commonly known under its brand name Herceptin, is a monoclonal anti-human epidermal growth factor 2 antibody that is used to treat HER2-positive cancers(24). It has an *N*-glycosylation site on position 297 of the heavy chain. One of the common glycans that occupy this glycosite is a monogalactosylated glycan with core-fucosylation (N4H4F1, F = fucose). As such, peptides from trastuzumab allowed us to test the effect of core fucosylation on the linkage variable. To this end, the protein was first digested by trypsin and subsequently split into two fractions. One of these fractions was defucosylated overnight, while both fractions were further partially digested with Pronase to obtain a range of glycopeptides of different sizes. LC-MS/MS analysis of these samples showed linkage variables ranging from 0.4 for EEQYNSTY with N4H4, to 0.58 for YNST with N4H4F1. Most glycan combinations yielded linkage variables in the 0.51-0.58 range, corresponding to an approximate 30-60% galactosylation of the 3-branch (**Figure 8B, C**). As far as we could see, core fucosylation had no apparent effect on the linkage variable, whereas the peptide appeared to have a minor effect. IgG N4H4(F1) is known to exist with both 3-and 6-branched galactosylation, although the ratio on trastuzumab in particular has not prior been reported. It is conceivable that interaction exists between galactose position and digestion efficiency of particular amino acids, but this would require substantial further investigation(25,26).

## DISCUSSION

While MS is nowadays one of the primary methods for glycan and glycoprotein analysis, its dependence on mass means that the important isomeric characteristics of glycosylation are difficult to access. However, by studying isomerically defined glycan standards, as well as sialylglycopeptide and trastuzumab, we here showed that several isomeric properties of glycosylation, namely sialic acid linkage and 3- vs. 6-linked branching, can be determined based on the relative intensities of glycan oxonium ions and even quantified in terms of relative ratios. By combining several oxonium ions across collision energies into a single “linkage variable”, we were able to discriminate isomer ratio differences as low as 10%, on a variety of peptides. Importantly, the MS scan time did not exceed 250 ms, bringing it in line with contemporary methods for glycoproteomics such as stepping-HCD and EThcD(8).

For now, the method was developed for precursors with a charge state of 2+, as these showed considerably larger differences in the fragmentation pattern across isomers when compared to precursors with 3+ charge. For the selection of product ions for the creation of the linkage variable only oxonium ions (B-ions) were included - this to minimize peptide- (and Y-ion)-specific masses that would make broad application of the method difficult. Correspondingly, we could observe a consistent linkage variable across varying peptide lengths of sialylglycopeptide and trastuzumab, although it should be noted that the glycopeptides used in our experiments were relatively small and did not include all amino acids. In addition, since the oxonium ions of single sialic acids were the main components in the differentiation of α2,3- and α2,6-linked sialylation and did not provide any significant improvements for branching isomerism, we excluded them from any of the linkage variables of the other glycopeptide isomers that contained sialic acids. By using the sialic acid oxonium ions only for the differentiation of α2,3- and α2,6-linked sialylation we hope to be able to combine branching and sialic acid linkage isomerism at a later date.

Interestingly, from our data it appears that the saccharides connected to the 3-branch of the glycan are generally fragmented at lower collision energies than the saccharide groups connected to the 6-branch of the glycan. A possible explanation for this consistent HCD fragmentation behavior is that linkages to the 3-carbon of a hexose are more rigid than linkages to the 6-carbon of a hexose. This seemed to be the case as well for the α2,3-linked NeuAc residues, which were more susceptible to dissociation than their α2,6-linked counterparts. It would be interesting to assess whether this is only the case for the structures we have investigated, or whether it proves to be a general rule that can be applied to future unknown glycopeptides. Currently, a main limiting factor for which isomeric properties can be analyzed is the availability of isomerically-defined glycosylated asparagine standards. General fragmentation rules, such as the 3-branch always fragmenting before the 6-branch, could help in distinguishing future unknown samples. However, for relative quantification glycopeptide, isomer mixtures still need to be defined and analyzed.

As it is, the method allows determination of the glycan structure of a glycopeptide without having peptide-specific parameters(16), apart from *m/z* ratio, intensity and retention time. In principle, this should enable easy incorporation into standard proteomics workflows, opening up a wide range of possibilities. For instance, it may be possible to generate a more detailed structural overview of antibody glycosylation in plasma samples, and thereby a better understanding of their differing effector functions and half-life characteristics(4). The method could also be instrumental in comparing normal human protein glycosylation to its alternative made by cell systems or even tumor cells – making the structural elements of glycosylation accessible would open up a new dimension of potential biomarker research(27).

For now, however, the method has not yet been tested in complex samples. More complex samples are expected to create a need for more complex data curation as well, and although the linkage variable currently allows for relative quantification of glycan structural elements, absolute quantification would require further investigation. On top of that, a complicating factor for system-wide glycoproteomics may be glycan-structure-specific proteolysis; it could very well be that certain proteoglycoforms are more easily cleaved by proteases than others, particularly when structural elements of glycosylation come into play. In addition, for purposes of biological investigation, it will be important to expand the repertoire of distinguishable glycan characteristics into directions such as α-and β-linkage-, fucose position and linkage, as well as antennary GlcNAc position. One way to achieve this may be the combination of our workflow with contemporary separation methods such as ion mobility.

As it is, we have developed a glycoproteomics-compatible method for distinguishing between glycopeptide isomers for branching position and sialic acid linkage. The method is largely independent of the connected peptide, and is expected to allow for the first instances of structural glycoproteomics by use of MS/MS alone.

## METHODS

### Chemicals and reagents

Milli-Q water was generated from Merck Milli-Q IQ 7003 system (Darmstadt, DE). Pronase and trastuzumab were obtained from Roche (Woerden, NL). ?-(1-2,3,4,6)-L-fucosidase was obtained from Megazyme (Ayr, UK). Chloroacetamide (CAA), Tris(2-carboxyethyl)phosphine hydrochloride (TCEP), Tris, trypsin, Lys-C, sodium deoxycholate (SDC), sodium acetate and formic acid (FA) were obtained from Merck (Darmstadt, DE). Acetonitrile and 0.1% FA were obtained from Biosolve (Valkenswaard, NL). Egg yolk powder was obtained by freeze-drying egg yolks from eggs obtained from SPAR (Waalwijk, NL). Cotton string was obtained from Kruidvat (Renswoude, NL). Trifluoric acid (TFA) was obtained from Honeywell International Inc (Charlotte, NC).

### Glycosylated asparagine standards

Isomerically-defined glycosylated asparagine standards were synthesized as previously described(21,22). The standards varied in either galactose positioning (3-branch and 6-branch), sialic acid positioning (3-branch and 6-branch, one galactose and two) or sialic acid linkage (α2,3- and α2,6-linked). Mixes of each isomer pair were made ranging from 100% isomer A and 0 % isomer B to 0% isomer A and 100% isomer B in steps of 10%.

### Glycopeptides digestion and purification

Sialylglycopeptide (SGP) and trastuzumab were used as model proteins for linkage determination. SGP was purified from egg yolk powder using HILIC-SPE. In short, 500 μg of egg yolk powder was dissolved in 250 μl 80% ACN 0.1% TFA. One cm of cotton string was used as column material. The column was conditioned with 3 times 200 μL 0.5% TFA in Milli-Q, then washed with 3 times 200 μL 80% ACN 0.5% TFA. After washing, the sample was loaded into the column 10 times. After sample loading the column was washed again with 3 times 200 μL 80% ACN 0.5% TFA and subsequently eluted with 2 times 200 μL 50% ACN 0.5% TFA. The eluent was dried in a vacuum concentrator and subsequently brought to 100 μL 0.1 M Tris 10 mM CaCl_2_ for Pronase digestion.

The Pronase digestion was performed at 37 °C and stopped at 7.5, 15, 30, 60, 120, 240 min, and after overnight digestion, to obtain a range of glycopeptides with varying peptide length. Trastuzumab was first reduced and alkylated in a pH 8.5 100 mM TRIS, 10 mM TCEP, 40 mM chloroacetamide 1% SDC buffer for 30 min at room temperature. Subsequently, the reduced and alkylated trastuzumab was digested overnight using trypsin and Lys-C in 50 mM ammonium bicarbonate at room temperature. Half of the tryptic trastuzumab digest was dissolved in 100 mM NaAc pH 4.5 and defucosylated using α-(1-2,3,4,6)-L-fucosidase overnight at 37 °C. Both the trastuzumab tryptic digest and the defucosylated trastuzumab were split into two fractions. One fraction was purified using the same HILIC-SPE protocol as the SGP, the other fraction was not further purified. All fraction were digested with Pronase for 15, 30, 60, 120 min and overnight.

### LC-MS method

The measurements were performed on Thermo scientific Exploris 480 and Fusion systems connected to Thermo Scientific Ultimate 3000 UHPLC systems. A 50 cm, 75 um ID, 2.4um reprosil column was used with a 60 min gradient of 2% B at 0-1 min, 13% B at 13 min, 44% B at 42 min, 99% B at 44-49 min, 2% B 50-60 min. After an initial MS scan, ions with a 2+ charge and an intensity of over 5·10^3^ were compared to a precursor list (*m/z* values: 127.0390, 138.0550, 145.0495, 163.0601, 168.0655, 186.0761, 204.0867, 243.0264, 274.0921, 292.1027, 366.1395, 405.0793, 407.1660, 512.1974 and 657.2349), which was used to trigger a MS/MS scan using higher-energy collisional dissociation (HCD) at 29% normalized collision energy (NCE). If at least 3 of aforementioned oxonium ions were detected, 10 subsequent MS/MS scans were performed of the same precursor using NCEs 5%-50%. To minimize any potential influence of the scan order on results, a semi-random scan order was used (NCEs: 30%, 10%, 50%, 20%, 40%, 15%, 45%, 25%, 35% and 5%) and the order was repeated in triplicate. After the MS/MS scans, the precursor was excluded from triggering further MS/MS scans for 15 s. To determine the optimal charge state, the method described above differed by not only covering 2+, but rather all charge states between 2+ and 8+.

### Data analysis

Thermo raw files were converted to MGF format using Proteowizard MSconvert (version 3.0.21328-404bcf1) using MGF as output format, 64-bit binary encoding precision and the following options selected: write index, zlib compression and TPP compatibility. No filters were used during the conversion from RAW files to MGF files. MGF files were searched for spectra containing glycan oxonium ions using an internally developed tool named PeakSuite (v1.10.1) using an ion delta of 20 ppm, noise filter of 0% and using a list of oxonium ion *m/z* values as mass targets (**Supplementary Table 1**). Among the scans, those without any detected peaks were removed, while those with oxonium ions were converted into text files. Python 3.2.2 was then used to 1) assign collision energies to the spectra, 2) for data curation based on retention time, precursor *m/z* and oxonium ion intensity and 3) for calculating the relative intensity of the oxonium ions (the Python script can be found as **Supplementary Information 1**). The relative intensities of the oxonium ions were calculated by dividing the intensity of an oxonium ion by the summed intensities of all oxonium ions in our oxonium ion list in the same scan.

The curated data was analyzed using RStudio (2022.02.2+485). For the glycosylated asparagine standards, levelplots were made for each of the aforementioned oxonium ions with the NCE on the x-axis (seq(5,50,1)), the percentage of isomer A on the y-axis (seq(0,100,2)) and relative intensity of the oxonium ions for the z-axis (the R script can be found as **Supplementary Information 2**). On basis of these outcomes, a variable was constructed that would report on the isomer ratios in each of the pair mixtures. For a full overview of linkage variable calculations, see **Supplementary Table 2**. Relevant variables were subsequently used to report on the isomeric properties of SGP and trastuzumab.

### Statistical information

To investigate differences in linkage variable for the mixtures of isomerically defined glycosylated asparagine standards, a two-tailed student’s t-test was performed between each two incremental mixtures (**Supplementary Table 3**). Results were deemed significant at p-value ≤ 0.05, with a false discovery rate of 5%.

### Data representation

To enhance the figures that were constructed with R, Microsoft Excel and Microsoft Powerpoint, glycan cartoons were generated using Glycoworkbench 2.1 following the recommendations of the Consortium for Functional Glycomics(28,29).

### Data availability

The data associated with the manuscript has been deposited on the MassIVE repository, and can be accessed at “ftp://massive.ucsd.edu/MSV000091172/” or by logging in with the username “MSV000091172_reviewer” and password “GlycoMSMS23”.

## ACKNOWLEDGEMENTS

We would like to thank Roche Diagnostics for kindly providing Pronase and tastuzumab, Dario Cramer for helpful initial digestion experiments, Hillary Hoppenbrouwers for her assistance with setting up cotton-HILIC-based SGP extraction from egg yolk powder, and Gerlof Bosman for his efforts on glycopeptide synthesis. Furthermore, we would like to thank Javier Sastre Toraño for providing the isomerically-defined glycosylated asparagine standards.

This project was funded by the Dutch Research Council (NWO) project VI.Veni.192.058, awarded to K.R.R. G.-J.P.H.B. acknowledges ERC Advanced Grant 101020769.

## AUTHOR CONTRIBUTIONS

J.C.L.M. performed experiments and data analysis and wrote the manuscript. G.-J.P.H.B. provided isomerically-defined glycosylated asparagine standards. J.M.A.D. performed experiments. K.R.R. performed experiments and data analysis, wrote the manuscript, designed the project and contributed the funding. All authors reviewed and edited the manuscript.

## MATERIALS & CORRESPONDENCE

Correspondence and material requests may be addressed to Karli R. Reiding

