## Supplementary material for "Glycoproteomics-compatible MS/MS-based quantification of glycopeptide isomers": Supplemantary Figures

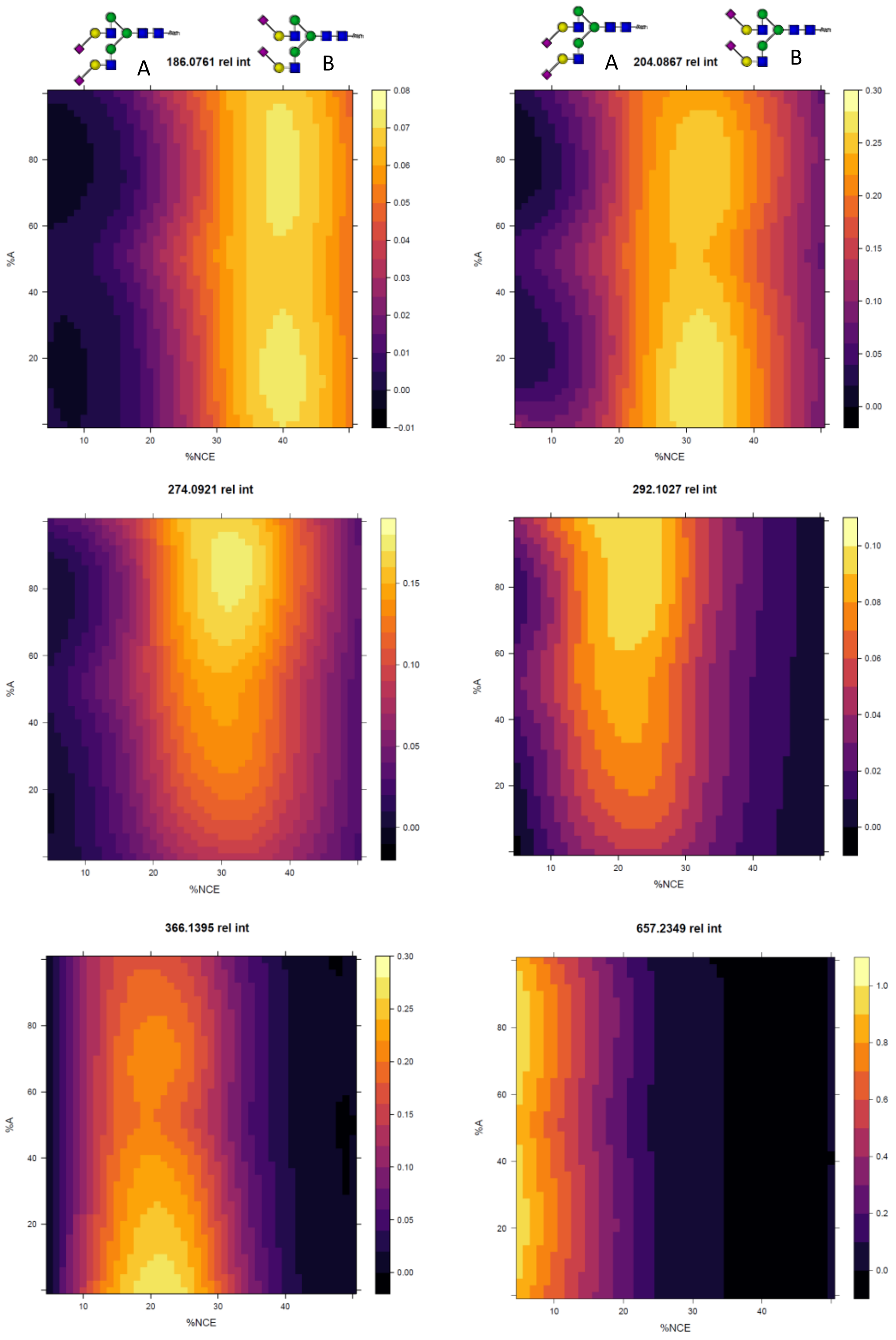


**Supplemental Figure 1. Relative intensities of oxonium ions for sialic acid linkage.** Relative intensities of oxonium ions are shown over a NCE range 5-50% and isomer mixtures ranging from 0-100% A. The color gradient indicates the relative intensity. HexNAc (*m/z* 204.0867), HexHexNAc (*m/z* 366.1395), NeuAc (*m/z* 292.1027) and NeuAc-H2O (*m/z* 274.0921) showed the biggest change between isomers.

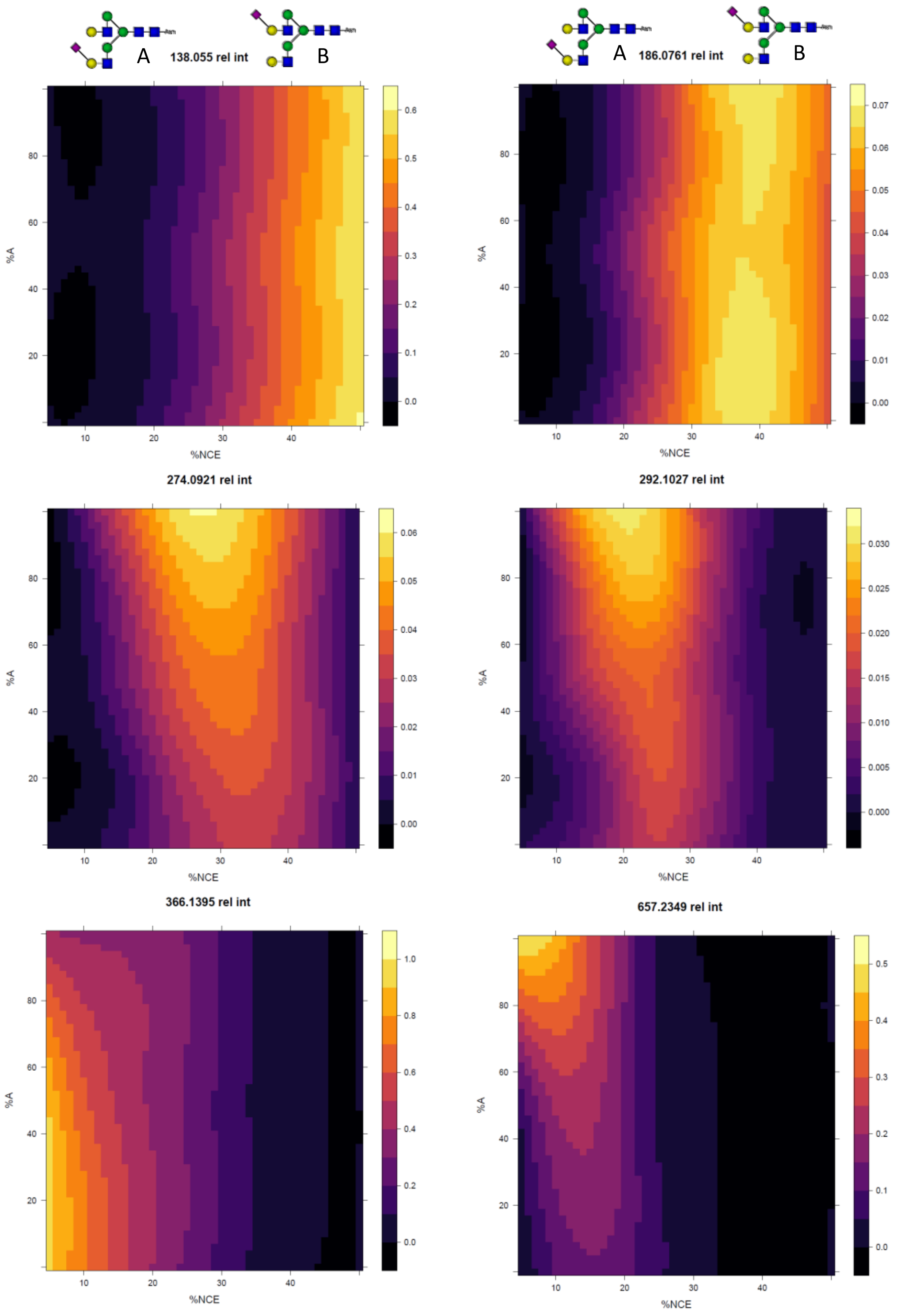


**Supplemental Figure 2. Relative intensities of oxonium ions for sialic acid branching.** Relative intensities of oxonium ions are shown over a NCE range 5-50% and isomer mixtures ranging from 0-100% A. The color gradient indicates the relative intensity. HexHexNAc (*m/z* 366.1395) and NeuAcHexHexNAc (*m/z* 657.2349) showed the biggest change between isomers.

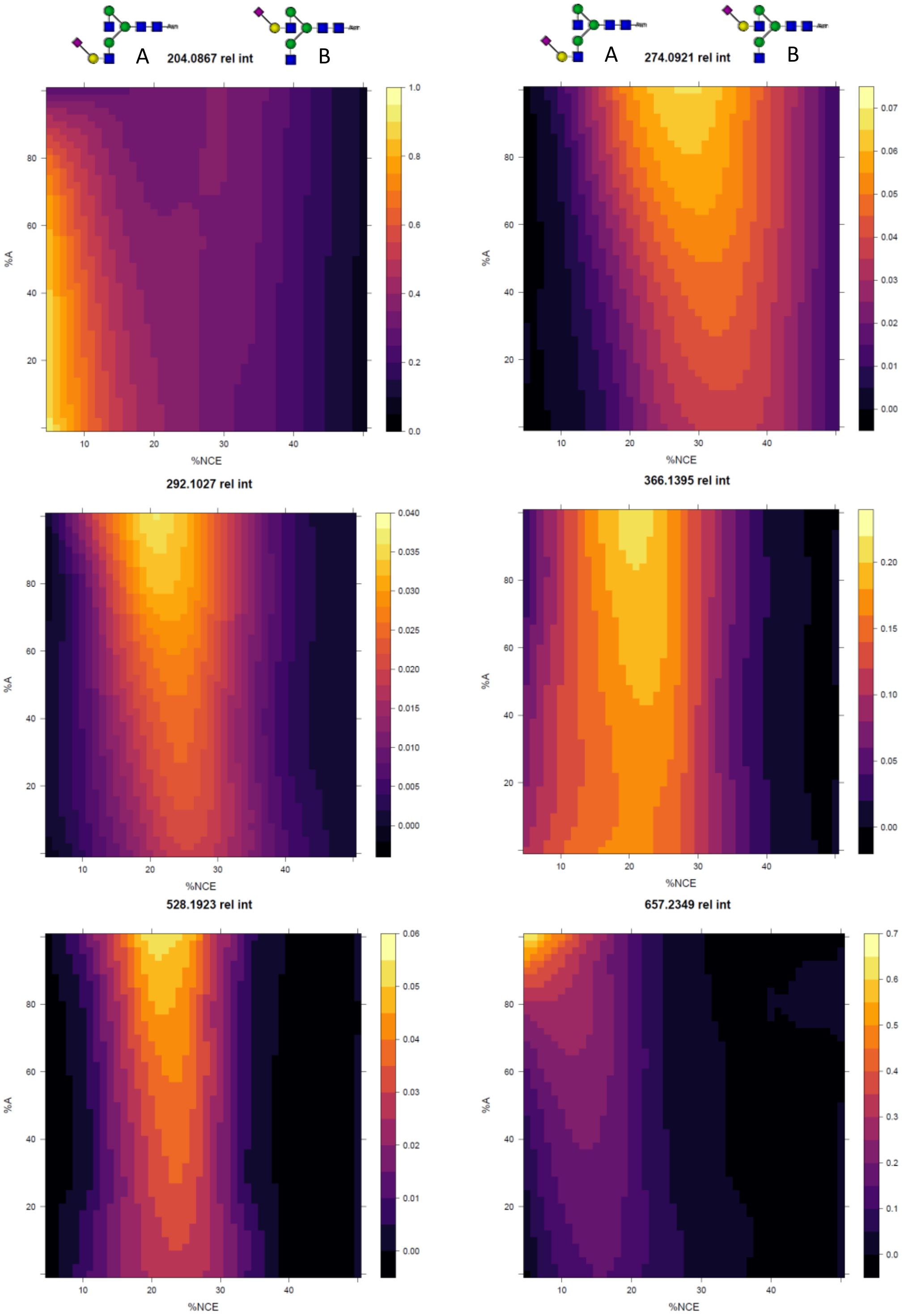


**Supplemental Figure 3. Relative intensities of oxonium ions for sialylated galactose branching.** Relative intensities of oxonium ions are shown over a NCE range 5-50% and isomer mixtures ranging from 0-100% A. The color gradient indicates the relative intensity.

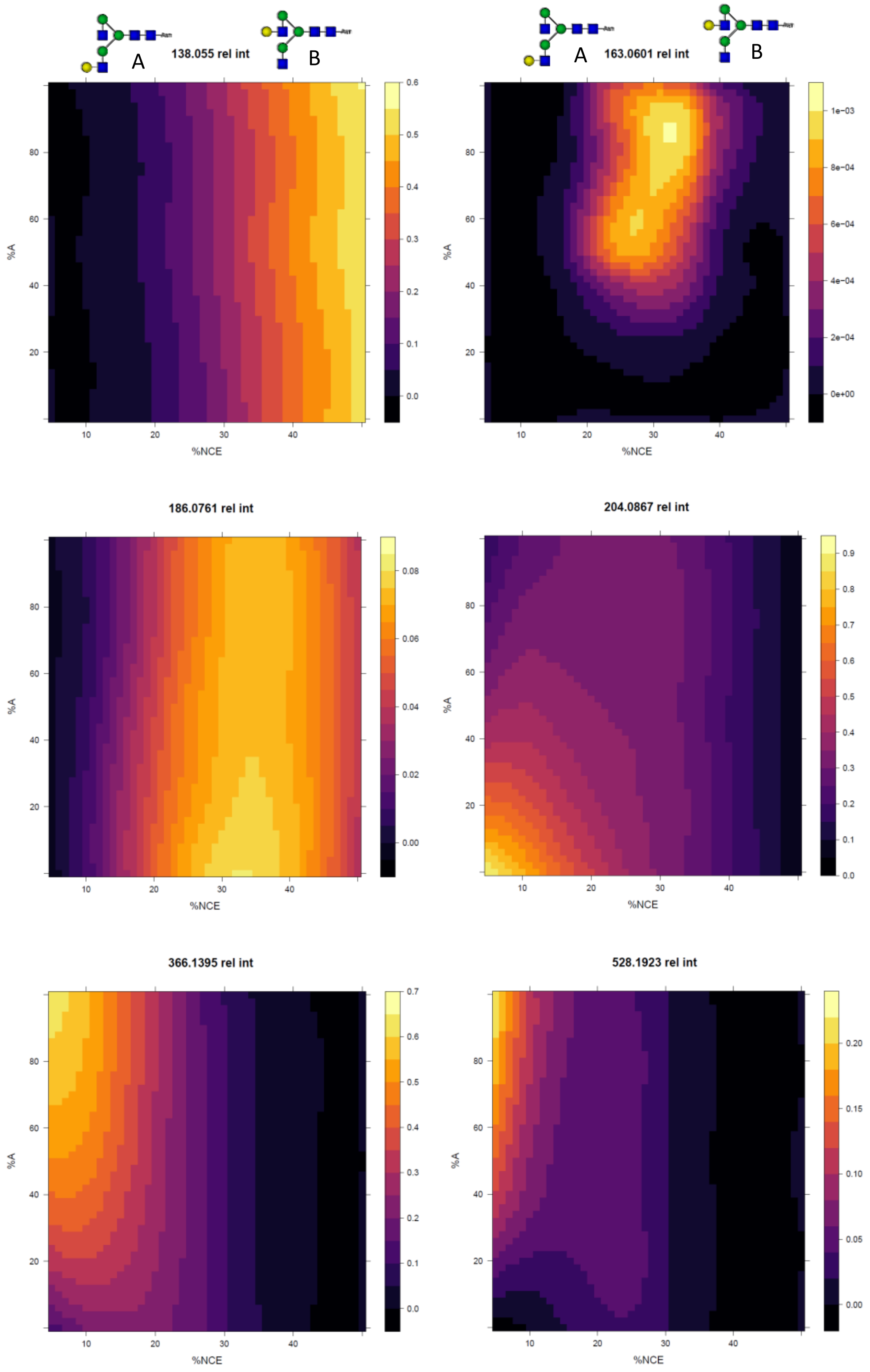


**Supplemental Figure 4. Relative intensities of oxonium ion for galactose branching.** Relative intensities of oxonium ions are shown over a NCE range 5-50% and isomer mixtures ranging from 0-100% A. The color gradient indicates the relative intensity. HexNAc (*m/z* 204.0867) and HexHexNAc (*m/z* 366.1395) showed the biggest change between isomers.

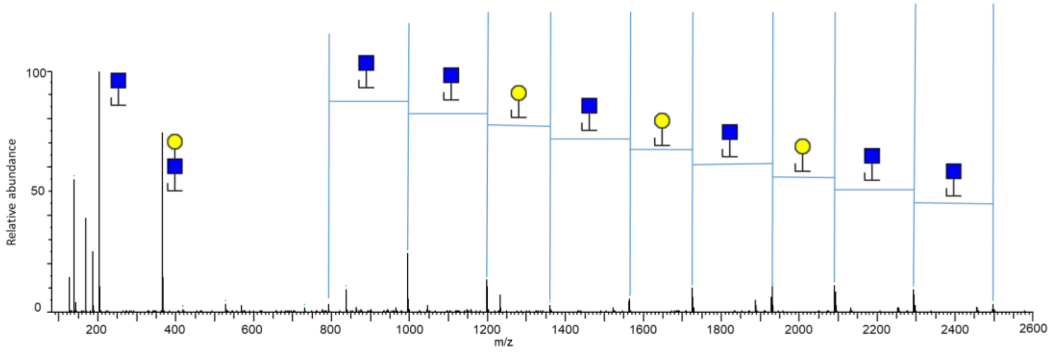


**Supplemental Figure 5: MS/MS spectrum showing an incorrect fragmentation pattern for SGP KVANKT-H4N5S2.** While having the same *m/z* value as KVANKT-H4N5S2 (*m/z* 1433.0890 at z=2+), the spectrum above clearly displays at least 6 HexNAc residues, suggesting a different glycan entirely. To avoid false assignments of glycopeptides such as these, an intensity threshold for certain oxonium ions was implemented, in this case a minimum of $1 \cdot 10^4$ for *m/z* 204.0867, 366.1395 and 657.2349 and $5 \cdot 10^3$ for *m/z* 274.0921 and 292.1027.

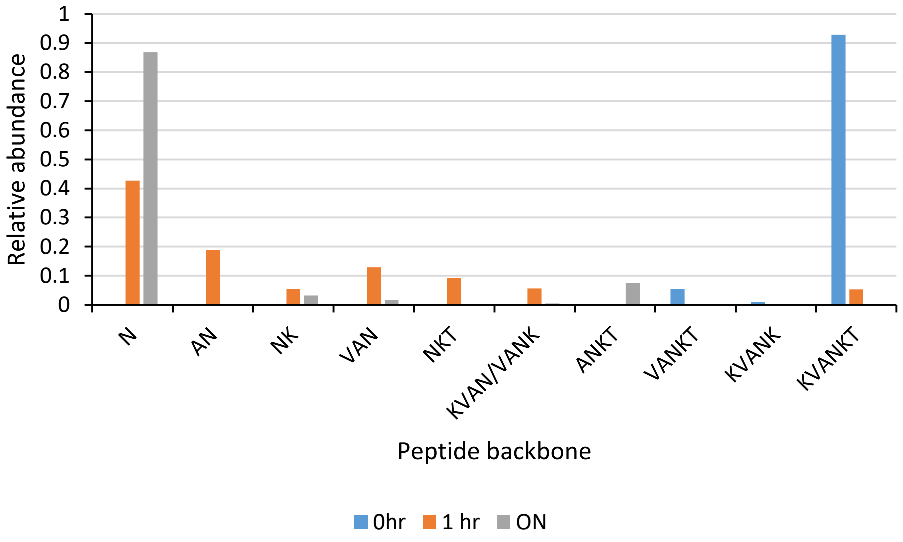


**Supplemental Figure 6: Partial proteolysis of SGP.** SGP was digested with Pronase for 1 hour or overnight. After 1 hour, already over 40% of the SGP was fully digested, with only a single asparagine still being connected to the glycan. After overnight digestion almost 90% of the SGP was completely digested. The partial digestion of SGP created a range of glycopeptide with the same glycan but varying peptide backbones.
