## Supplementary Information 1 for "Glycoproteomics-compatible MS/MS-based quantification of glycopeptide isomers"

#! /usr/bin/env python

##########################################

\### Glycopeptide ismer data analysis###

\# Libraries

import os

import sys

import re

import pandas as pd

from time import perf_counter

\### Functions ####

def _LVCombine(dfs,LVName,NCEList):

'''Generate LV from combined dfs lines.'''

LV = 0.0

for NCE in NCEList:

LVs = dfs[dfs.index == NCE][LVName]

if LVs.isnull().any():

LVs.iat[0] = 0.0

try:

LV += LVs.iat[0]

except IndexError:

LV += 0.0

LV = LV / len(NCEList)

return(LV)

\### Main loop ###

def main():

'''Main loop.'''

\### Timing

t1 = perf_counter()

\### Input list control

removeList = ["Unnamed","mz="," delta"," int/basepeak",

"scan number","id"]

cycleName = "NCE"

\### NCE cycle and removelists

NCEcycle_single = ["29","30","10","50","20","40",

"15","45","25","35","05"]

NCEcycle = NCEcycle_single*3

removeDictNCE = {}

for i in NCEcycle:

removeDictNCE[i] = NCEcycle[:]

removeDictNCE[i].remove(i)

\### Ion m/z values - this could be read from file

ionList = [ "126.055" , "127.039" , "138.055" , "144.0655",

"145.0495", "163.0601", "168.0655", "186.0761",

"204.0867", "243.0264", "274.0921", "290.087" ,

"292.1027", "308.0976", "366.1395", "405.0793",

"407.166" , "454.1555", "512.1974", "528.1923",

"569.2188", "657.2349", "690.2451", "731.2717",

"819.2877", "852.2979", "893.3245", "934.351" ,

"981.3405","1055.3773","1096.4039","1143.3934",

"1184.4199","1217.4301","1258.4567","1299.4832",

"1346.4727","1420.5095","1461.5361","1508.5256",

"1549.5521","1582.5623","1623.5889","1711.6049",

"1785.6417","1799.621" ,"2002.7003"]

ionIList = []

for i in ionList:

ionIList.append(i + " int")

\### Generating columns list from ion list

columnList =["title","A","B","NCE","retention time",

"precursor mz","precursor int","precursor charge"]

for i in ionList:

columnList.append(i + " int")

columnList = columnList + ["ion sum"]

for i in ionList:

columnList.append(i + " rel int")

columnList = columnList + ["int_time","linkage variable",

"sialic sum","hexnac sum",

"glycan check"]

\### building block masses

AADict = {"G": 57.02146,

"A": 71.03711,

"S": 87.03203,

"P": 97.05276,

"V": 99.06841,

"T":101.04768,

"C":103.00919,

"I":113.08406,

"L":113.08406,

"N":114.04293,

"D":115.02694,

"Q":128.05858,

"K":128.09496,

"E":129.04259,

"M":131.04049,

"H":137.05891,

"F":147.06841,

"R":156.10111,

"Y":163.06333,

"W":186.07931}

ModDict = {"P": 79.96633,

"F":146.05791,

"H":162.05282,

"N":203.07937,

"S":291.09542,

"A":291.09542,

"G":307.09033,

"Carbamidomethylation": 57.02146,

"Water": 18.01057,

"Proton":1.00728,

"Electron":0.00055}

\### Input potential glycopeptides

gpList = ["EEQYNSTYR	N4H4",

"EEQYNSTYR	N4H4F1",

"EEQYNSTY	N4H4F1",

"NSTY	N4H4F1",

"EEQFNSTYR	N4H4",

"EEQFNSTYR	N4H4F1",

"EEQFNSTY	N4H4F1",

"NSTY	N4H4F1",

"EEQFNSTFR	N4H4",

"EEQFNSTFR	N4H4F1",

"EEQFNSTF	N4H4F1",

"NSTF	N4H4F1",

"EQYN	N4H4F1",

"EQYN	N4H4",

"YNST	N4H4F1",

"YNST	N4H4",

"EEQYN	N4H4F1",

"EEQYN	N4H4",

"EEQYNST	N4H4F1",

"EEQYNST	N4H4",

"EQYNST	N4H4F1",

"EQYNST	N4H4",

"TKPREEQYNSTYR	N4H4",

"TKPREEQYNSTYR	N4H4F1",

"TKPREEQYNSTY	N4H4F1",

"TKPREEQFNSTYR	N4H4",

"TKPREEQFNSTYR	N4H4F1",

"TKPREEQFNSTY	N4H4F1",

"TKPREEQFNSTFR	N4H4",

"TKPREEQFNSTFR	N4H4F1",

"TKPREEQFNSTF	N4H4F1",

"LREEQFNSTFR	N4H4",

"LREEQFNSTFR	N4H4F1",

"LREEQFNSTF	N4H4F1",

"TKLREEQFNSTFR	N4H4",

"TKLREEQFNSTFR	N4H4F1",

"TKLREEQFNSTF	N4H4F1",

"TKPWEEQFNSTFR	N4H4",

"TKPWEEQFNSTFR	N4H4F1",

"TKPWEEQFNSTFR	N4H4F1"]

for i in range(len(gpList)):

p,g = gpList[i].split()

gpList[i] = p + " " + g

\### Generating and storing glycopeptide masses

ppm = 10	# ppm tolerance for m/z values - was 5, maybe 10?

ppm_MS1 = 10

ppm_MS2 = 20

zList = [2]	# In case of multiple charges

gpDict = {}

for gp in gpList:

p,g = gp.split()

for z in zList:

zStr = str(z)

zStrL = zStr + "_lower"

zStrU = zStr + "_upper"

gpDict[gp] = {"peptide":p,

"glycan":g,

zStr:0,

zStrL:0,

zStrU:0}

\# Calculate masses

for aa in p:

gpDict[gp][zStr] += AADict[aa]

gList = re.split("(\d+)",g)

for i in range(len(gList)):

if gList[i].isdigit():

continue

elif not gList[i] == "":

gpDict[gp][zStr] += (ModDict[gList[i]] *

float(gList[i+1]))

gpDict[gp][zStr] += ModDict["Water"]

gpDict[gp][zStr] += ModDict["Proton"]*z

gpDict[gp][zStr] = round(gpDict[gp][zStr]/z,5)

gpDict[gp][zStrL] = round(gpDict[gp][zStr] -

(ppm / 1e6 * gpDict[gp][zStr]),5)

gpDict[gp][zStrU] = round(gpDict[gp][zStr] +

(ppm / 1e6 * gpDict[gp][zStr]),5)

\### Making list of files

wd = "insertwdhere"

if wd == "":

wd = os.path.abspath("")

os.chdir(wd)

fList = []

for root, dirs, files in os.walk(".", topdown = False):

for name in files:

if ".txt" in name:

fList.append(name)

\### Reformatting and storing files within dataframe

print("Reading files...")

df = pd.DataFrame()

for f in fList:

print("...",f)

d = pd.read_table(f,sep="\t",header=0)

f = f.split("\\")[-1]

f = f.split(".")[0]

d["filename"] = f

\# Prevent issue when changing from 9.99 to 10.00 min RT

d["rt_min"] = d["retention time"] / 60

d["rt_minx10000"] = round(d["rt_min"]*1e4,0)

d["rt_minx10000"] = d["rt_minx10000"].astype(str)

d["rt_minx10000"] = d["rt_minx10000"].str.zfill(12)

\# Build and sort on unique rt+int values

d["int_time"] = d["precursor int"].astype(str)+d["rt_minx10000"]

d.sort_values("int_time",inplace=True)

\# Assign NCE values from NCEcycle

#print("NCE cycle assignment...")

cList = []

int_prev = 0

for int_data in d["precursor mz"]:

if int_data == int_prev:

cList.append(NCEcycle[NCEcycle.index(cList[-1])+1])

else:

cList.append(NCEcycle[0])

int_prev = int_data

d[cycleName] = cList[:] #recycle script change

\# Calculate relative intensities

#print("Relative intensity calculation...")

d[ionIList]= d[ionIList].replace("-","0")

d[ionIList]= d[ionIList].apply(pd.to_numeric,errors="coerce")

d["ion sum"] = d[ionIList].sum(axis=1)

for ion in ionList:

d[ion + " rel int"] = d[ion + " int"] / d["ion sum"]

\# Peptide assignment

#print("m/z-based glycopeptide assignment...")

d["A"] = "-"

d["Sequence"] = "-"

d["Glycan"] = "-"

for gp in gpDict:

p,g = gp.split()

for z in zList:

gpzL = gpDict[gp][str(z)+"_lower"]

gpzU = gpDict[gp][str(z)+"_upper"]

d.loc[((d["precursor mz"] >= gpzL) &

(d["precursor mz"] <= gpzU)),

"A"] = gp

d.loc[((d["precursor mz"] >= gpzL) &

(d["precursor mz"] <= gpzU)),

"sequence"] = p

d.loc[((d["precursor mz"] >= gpzL) &

(d["precursor mz"] <= gpzU)),

"glycan"] = g

\# Preliminary linkage variable calculations

#print("Linkage variable calculations...")

d["LV_3v6-branch Gal"]	= (d["204.0867 rel int"] /

(d["204.0867 rel int"] +

d["366.1395 rel int"]))

d["LV_3v6-branch SiaGal"]	= (d["204.0867 rel int"] /

(d["204.0867 rel int"] +

d["366.1395 rel int"] +

d["657.2349 rel int"]))

d["LV_3v6-branch Sia"]	= (d["366.1395 rel int"] /

(d["366.1395 rel int"] +

d["657.2349 rel int"]))

d["LV_3v6-linked Sia"]	= ((d["274.0921 rel int"] +

d["292.1027 rel int"])/

(d["274.0921 rel int"] +

d["292.1027 rel int"] +

d["204.0867 rel int"] +

d["366.1395 rel int"]))

\# Add file to dataframe

df = df.append(d)

\### Data quality control

print("Data quality control...")

\## Intensity threshold QC

IntQC_204 = 1e4

IntQC_274 = 5e3

IntQC_292 = 5e3

IntQC_366 = 1e4

IntQC_657 = 1e4

\# 204

df.loc[((df["204.0867 int"] >= IntQC_204) &

(df["NCE"] == "30")),"QC_204"] = "pass"

df.loc[((df["204.0867 int"] < IntQC_204) &

(df["NCE"] == "30")),"QC_204"] = "fail"

\# 274

df.loc[((df["274.0921 int"] >= IntQC_274) &

(df["NCE"] == "30")),"QC_274"] = "pass"

df.loc[((df["274.0921 int"] < IntQC_274) &

(df["NCE"] == "30")),"QC_274"] = "fail"

\# 292

df.loc[((df["292.1027 int"] >= IntQC_292) &

(df["NCE"] == "30")),"QC_292"] = "pass"

df.loc[((df["292.1027 int"] < IntQC_292) &

(df["NCE"] == "30")),"QC_292"] = "fail"

\# 366

df.loc[((df["366.1395 int"] >= IntQC_366) &

(df["NCE"] == "30")),"QC_366"] = "pass"

df.loc[((df["366.1395 int"] < IntQC_366) &

(df["NCE"] == "30")),"QC_366"] = "fail"

\# 657

df.loc[((df["657.2349 int"] >= IntQC_657) &

(df["NCE"] == "30")),"QC_657"] = "pass"

df.loc[((df["657.2349 int"] < IntQC_657) &

(df["NCE"] == "30")),"QC_657"] = "fail"

\# Fill in lists

for QC in ["QC_204","QC_274","QC_292","QC_366","QC_657"]:

df.loc[:,QC]=df.loc[:,QC].ffill()

\## RT window QC

rtQC_lower = 10.0	# Should be set per analyte - via file?

rtQC_upper = 20.0	# Stored in gpDict?

df["QC_rt"] = "fail"

for gp in gpDict:

df.loc[((df["A"] == gp) &

(df["rt_min"] >= rtQC_lower) &

(df["rt_min"] <= rtQC_upper)),

"QC_rt"] = "pass"

\### Removing and reorganizing columns

print("Removing unnecessary columns...")

\# Removal

for key in df.keys():

for rm in removeList:

if rm in key:

df.drop(key, axis=1, inplace=True)

\### Write to file

print("Writing complete list to file...")

print("... csv")

df.set_index("NCE", inplace=True)

df.to_csv(wd + "insertnamehere.csv")

#print("... xlsx") # Slow compared to csv

#writer = pd.ExcelWriter(wd + "\\_Combined.xlsx",

\#	engine = "xlsxwriter",

\#	options={"strings_to_numbers": True})

#df.to_excel(writer, sheet_name = "all data")

#writer.save()

#writer.close()

"""

\### Timing report

t2 = perf_counter()

print("Done! Elapsed time:",round(t2-t1,4),"seconds.")

return(0)

"""

\### Generate new dataset with one peptide per row

print("Calculating precursor-specific linkage variables...")

dfu = pd.DataFrame(columns = df.columns) # df unique

uniqueList = df["precursor int"].unique()

for unique in uniqueList:

dfs = df[df["precursor int"] == unique] # df sub

\## Inherit properties from NCE 29 scan

dfs_29 = dfs[dfs.index == "30"]

\## Linkage variable combinations

\# 3v6-branch Gal	(10-25%)

LVName = "LV_3v6-branch Gal"

NCEList = ["10","15","20","25"]

dfs_29.loc["30",LVName] = _LVCombine(dfs,LVName,NCEList)

\# 3v6-branch SiaGal	(10-20%)

LVName = "LV_3v6-branch SiaGal"

NCEList = ["10","15","20"]

dfs_29.loc["30",LVName] = _LVCombine(dfs,LVName,NCEList)

\# 3v6-branch Sia	(10-15%)

LVName = "LV_3v6-branch Sia"

NCEList = ["10","15"]

dfs_29.loc["30",LVName] = _LVCombine(dfs,LVName,NCEList)

\# 3v6-linked Sia	(20-45%)

LVName = "LV_3v6-linked Sia"

NCEList = ["20","25","30","35","40","45"]

dfs_29.loc["30",LVName] = _LVCombine(dfs,LVName,NCEList)

\# Assign and build new dataframe

dfu = dfu.append(dfs_29,ignore_index=False)

\### Write new dataset to file

print("Writing unique precursors to file...")

print("... csv")

dfu.to_csv(wd + "\\insertnamehere.csv")

\### Timing report

t2 = perf_counter()

print("Done! Elapsed time:",round(t2-t1,4),"seconds.")

return(0)

if __name__ == "__main__":

main()

sys.exit()
