## Supplementary Information 2 for "Glycoproteomics-compatible MS/MS-based quantification of glycopeptide isomers"

\###### Script for glycopeptide isomer data analysis ######

####### Joshua Maliepaard 30-1-2023 #######

\### Installs ###

#install.packages("plot3D")

#install.packages("lattice")

#install.packages("viridis")

#install.packages("tidyverse")

\### Imports ###

library(plot3D)

library(lattice)

library(viridis)

library(tidyverse)

\### Variables ###

fn_input = "Insertfilename.txt"###input file

wd = "insertworkingdirectory"###working directory

subset_start = 67### first column with oxonium ion relative intensities.

subset_end = 113### Last column with oxonium ion relative intensities.

\### Init ###

setwd(wd)

dat = read.table(fn_input,sep=",",header=1,na.string=c(NA,""),fill=T, check.names=FALSE)

############################################################

\### Boxplots, scatterplots and levelplots of oxonium ions###

############################################################

\### Boxplots

list_z = names(dat)[subset_start:subset_end]

list_A = levels(factor(dat$X.A.))

pdf(file="nameoutputfile.pdf")

par(mfrow=c(3,4))

for (var_z in list_z){

boxplot(dat[,var_z]~dat$NCE,na.rm=T,

ylim=c(0,1),

xlab="%NCE",ylab="RI",main=var_z)

for (A in list_A){

dat_sub = dat[which(dat$X.A.==A),]

boxplot(dat_sub[,var_z]~dat_sub$NCE,na.rm=T,

ylim=c(0,1),

xlab="%NCE",ylab="RI",main=paste(var_z,A))

}

}

dev.off()

\### Scatterplots

list_z = names(dat)[subset_start:subset_end]

list_A = levels(factor(dat$X.A.))

pdf(file="nameoutputfile.pdf")

par(mfrow=c(3,4))

for (var_z in list_z){

plot(dat[,var_z]~dat$NCE,na.rm=T,

ylim=c(0,1),

xlab="%NCE",ylab="RI",main=var_z)

for (A in list_A){

dat_sub = dat[which(dat$A==A),]

plot(dat_sub[,var_z]~dat_sub$NCE,na.rm=T,

ylim=c(0,1),

xlab="%NCE",ylab="RI",main=paste(var_z,A))

}

}

dev.off()

\### Level plot

list_z = names(dat)[subset_start:subset_end]

pdf(file="nameoutputfile.pdf")

for(var_z in list_z){

df = data.frame(x=dat$NCE,y=dat$A,z=dat[,var_z])

df.loess = loess(z~x*y,data=df,degree=2,span=0.25)

df.fit = expand.grid(list(x = seq(5,50,1), y = seq(0,100,2)))

z = predict(df.loess, newdata = df.fit)

height = as.numeric(z)

height[which(height<0)]=0

print(height)

z_plot = levelplot(height ~ x*y, data = df.fit,

xlab = "%NCE", ylab = "%A",

main = var_z,

col.regions = inferno(100),

pretty = T

)

print(z_plot)

}

dev.off()

#####################################

\### Linkage varaible calculations ###

#####################################

###Linkage variable oxonium ion for galactose branching

#form_nom = c("204.0867 rel int")### nominator for linkage variable

#form_dnom = c("204.0867 rel int","366.1395 rel int")### denominator for linkage variable

###Linkage variable oxonium ion for sialylated galactose branching

#form_nom = c("204.0867 rel int")### nominator for linkage variable

#form_dnom = c("204.0867 rel int","366.1395 rel int","657.2349 rel int")### denominator for linkage variable

###Linkage variable oxonium ion for sialic acid branching

#form_nom = c("366.1395 rel int")### nominator for linkage variable

#form_dnom = c("657.2349 rel int","366.1395 rel int")### denominator for linkage variable

###Linkage variable oxonium ions for sialic acid linkage

form_nom = c("292.1027 rel int", "274.0921 rel int")### nominator for linkage variable

form_dnom = c("292.1027 rel int","366.1395 rel int","274.0921 rel int","204.0867 rel int")### denominator for linkage variable

dat_NCE5 = dat[which(dat$NCE==5),]

dat_NCE10 = dat[which(dat$NCE==10),]

dat_NCE15 = dat[which(dat$NCE==15),]

dat_NCE20 = dat[which(dat$NCE==20),]

dat_NCE25 = dat[which(dat$NCE==25),]

dat_NCE30 = dat[which(dat$NCE==30),]

dat_NCE35 = dat[which(dat$NCE==35),]

dat_NCE40 = dat[which(dat$NCE==40),]

dat_NCE45 = dat[which(dat$NCE==45),]

dat_NCE50 = dat[which(dat$NCE==50),]

ms2_list = levels(factor(dat$"precursor int"))

ms2 = matrix(ms2_list)

ms2$int = ms2_list

ms2$A = array(dim=length(ms2_list))

ms2$NCE5 = array(dim=length(ms2_list))

ms2$NCE10 = array(dim=length(ms2_list))

ms2$NCE15 = array(dim=length(ms2_list))

ms2$NCE20 = array(dim=length(ms2_list))

ms2$NCE25 = array(dim=length(ms2_list))

ms2$NCE30 = array(dim=length(ms2_list))

ms2$NCE35 = array(dim=length(ms2_list))

ms2$NCE40 = array(dim=length(ms2_list))

ms2$NCE45 = array(dim=length(ms2_list))

ms2$NCE50 = array(dim=length(ms2_list))

for (int in ms2$int){

ms2$A[which(ms2$int == int)] = dat$"A"[which(dat$"precursor int" == int)][1]

}

for (int in ms2$int){

nom = dnom = 0

for (num in form_nom){nom = nom + dat_NCE5[which(dat_NCE5$"precursor int" == int),num]}

for (num in form_dnom){dnom = dnom + dat_NCE5[which(dat_NCE5$"precursor int" == int),num]}

if (length(nom/dnom) > 0){ms2$NCE5[which(ms2$int == int)] = nom/dnom}

}

for (int in ms2$int){

nom = dnom = 0

for (num in form_nom){nom = nom + dat_NCE10[which(dat_NCE10$"precursor int" == int),num]}

for (num in form_dnom){dnom = dnom + dat_NCE10[which(dat_NCE10$"precursor int" == int),num]}

if (length(nom/dnom) > 0){ms2$NCE10[which(ms2$int == int)] = nom/dnom}

}

for (int in ms2$int){

nom = dnom = 0

for (num in form_nom){nom = nom + dat_NCE15[which(dat_NCE15$"precursor int" == int),num]}

for (num in form_dnom){dnom = dnom + dat_NCE15[which(dat_NCE15$"precursor int" == int),num]}

if (length(nom/dnom) > 0){ms2$NCE15[which(ms2$int == int)] = nom/dnom}

}

for (int in ms2$int){

nom = dnom = 0

for (num in form_nom){nom = nom + dat_NCE20[which(dat_NCE20$"precursor int" == int),num]}

for (num in form_dnom){dnom = dnom + dat_NCE20[which(dat_NCE20$"precursor int" == int),num]}

if (length(nom/dnom) > 0){ms2$NCE20[which(ms2$int == int)] = nom/dnom}

}

for (int in ms2$int){

nom = dnom = 0

for (num in form_nom){nom = nom + dat_NCE25[which(dat_NCE25$"precursor int" == int),num]}

for (num in form_dnom){dnom = dnom + dat_NCE25[which(dat_NCE25$"precursor int" == int),num]}

if (length(nom/dnom) > 0){ms2$NCE25[which(ms2$int == int)] = nom/dnom}

}

for (int in ms2$int){

nom = dnom = 0

for (num in form_nom){nom = nom + dat_NCE30[which(dat_NCE30$"precursor int" == int),num]}

for (num in form_dnom){dnom = dnom + dat_NCE30[which(dat_NCE30$"precursor int" == int),num]}

if (length(nom/dnom) > 0){ms2$NCE30[which(ms2$int == int)] = nom/dnom}

}

for (int in ms2$int){

nom = dnom = 0

for (num in form_nom){nom = nom + dat_NCE35[which(dat_NCE35$"precursor int" == int),num]}

for (num in form_dnom){dnom = dnom + dat_NCE35[which(dat_NCE35$"precursor int" == int),num]}

if (length(nom/dnom) > 0){ms2$NCE35[which(ms2$int == int)] = nom/dnom}

}

for (int in ms2$int){

nom = dnom = 0

for (num in form_nom){nom = nom + dat_NCE40[which(dat_NCE40$"precursor int" == int),num]}

for (num in form_dnom){dnom = dnom + dat_NCE40[which(dat_NCE40$"precursor int" == int),num]}

if (length(nom/dnom) > 0){ms2$NCE40[which(ms2$int == int)] = nom/dnom}

}

for (int in ms2$int){

nom = dnom = 0

for (num in form_nom){nom = nom + dat_NCE45[which(dat_NCE45$"precursor int" == int),num]}

for (num in form_dnom){dnom = dnom + dat_NCE45[which(dat_NCE45$"precursor int" == int),num]}

if (length(nom/dnom) > 0){ms2$NCE45[which(ms2$int == int)] = nom/dnom}

}

for (int in ms2$int){

nom = dnom = 0

for (num in form_nom){nom = nom + dat_NCE50[which(dat_NCE50$"precursor int" == int),num]}

for (num in form_dnom){dnom = dnom + dat_NCE50[which(dat_NCE50$"precursor int" == int),num]}

if (length(nom/dnom) > 0){ms2$NCE50[which(ms2$int == int)] = nom/dnom}

}

###replace NA values with 0

ms2$NCE5 = replace(ms2$NCE5,is.na(ms2$NCE5),0)

ms2$NCE10 = replace(ms2$NCE10,is.na(ms2$NCE10),0)

ms2$NCE15 = replace(ms2$NCE15,is.na(ms2$NCE15),0)

ms2$NCE20 = replace(ms2$NCE20,is.na(ms2$NCE20),0)

ms2$NCE25 = replace(ms2$NCE25,is.na(ms2$NCE25),0)

ms2$NCE30 = replace(ms2$NCE30,is.na(ms2$NCE30),0)

ms2$NCE35 = replace(ms2$NCE35,is.na(ms2$NCE35),0)

ms2$NCE40 = replace(ms2$NCE40,is.na(ms2$NCE40),0)

ms2$NCE45 = replace(ms2$NCE45,is.na(ms2$NCE45),0)

ms2$NCE50 = replace(ms2$NCE50,is.na(ms2$NCE50),0)

###Linkage variable NCE range for galactose branching

#ms2$NCEsum = (ms2$NCE10 + ms2$NCE15 + ms2$NCE20 + ms2$NCE25)/4

###Linkage variable NCE range for sialylated galactose branching

#ms2$NCEsum = (ms2$NCE10 + ms2$NCE15 + ms2$NCE20)/3

###Linkage variable NCE range for sialic acid branching

#ms2$NCEsum = (ms2$NCE10 + ms2$NCE15)/2

###Linkage variable NCE range for sialic acid linkage

ms2$NCEsum = (ms2$NCE20 + ms2$NCE25 + ms2$NCE30 + ms2$NCE35 + ms2$NCE40 + ms2$NCE45)/6

\### Place NA values back

ms2$NCEsum = replace(ms2$NCEsum,ms2$NCEsum <= 0,NA)

ms2$NCE5 = replace(ms2$NCE5,ms2$NCE5 <= 0,NA)

ms2$NCE10 = replace(ms2$NCE10,ms2$NCE10 <= 0,NA)

ms2$NCE15 = replace(ms2$NCE15,ms2$NCE15 <= 0,NA)

ms2$NCE20 = replace(ms2$NCE20,ms2$NCE20 <= 0,NA)

ms2$NCE25 = replace(ms2$NCE25,ms2$NCE25 <= 0,NA)

ms2$NCE30 = replace(ms2$NCE30,ms2$NCE30 <= 0,NA)

ms2$NCE35 = replace(ms2$NCE35,ms2$NCE35 <= 0,NA)

ms2$NCE40 = replace(ms2$NCE40,ms2$NCE40 <= 0,NA)

ms2$NCE45 = replace(ms2$NCE45,ms2$NCE45 <= 0,NA)

ms2$NCE50 = replace(ms2$NCE50,ms2$NCE50 <= 0,NA)

################################

\### Plotting linkage variable###

################################

x = ms2$NCEsum

y = ms2$A

model = lm(x~y)

boxplot(x~y,xlab="%A",ylab="Linkage variable", ylim=c(0,1))

#abline(model)

#summary(model)

###############

\### t.tests ###

###############

set1 = ms2$NCEsum[which(ms2$A == "0")]

set2 = ms2$NCEsum[which(ms2$A == "10")]

t.test(set1,set2)

set1 = ms2$NCEsum[which(ms2$A == "10")]

set2 = ms2$NCEsum[which(ms2$A == "20")]

t.test(set1,set2)

set1 = ms2$NCEsum[which(ms2$A == "20")]

set2 = ms2$NCEsum[which(ms2$A == "30")]

t.test(set1,set2)

set1 = ms2$NCEsum[which(ms2$A == "30")]

set2 = ms2$NCEsum[which(ms2$A == "40")]

t.test(set1,set2)

set1 = ms2$NCEsum[which(ms2$A == "40")]

set2 = ms2$NCEsum[which(ms2$A == "50")]

t.test(set1,set2)

set1 = ms2$NCEsum[which(ms2$A == "50")]

set2 = ms2$NCEsum[which(ms2$A == "60")]

t.test(set1,set2)

set1 = ms2$NCEsum[which(ms2$A == "60")]

set2 = ms2$NCEsum[which(ms2$A == "70")]

t.test(set1,set2)

set1 = ms2$NCEsum[which(ms2$A == "70")]

set2 = ms2$NCEsum[which(ms2$A == "80")]

t.test(set1,set2)

set1 = ms2$NCEsum[which(ms2$A == "80")]

set2 = ms2$NCEsum[which(ms2$A == "90")]

t.test(set1,set2)

set1 = ms2$NCEsum[which(ms2$A == "90")]

set2 = ms2$NCEsum[which(ms2$A == "100")]

t.test(set1,set2)
